# Kiwa is a bacterial membrane-embedded defence supercomplex activated by phage-induced membrane changes

**DOI:** 10.1101/2023.02.26.530102

**Authors:** Zhiying Zhang, Thomas C. Todeschini, Yi Wu, Roman Kogay, Ameena Naji, Joaquin Cardenas Rodriguez, Rupavidhya Mondi, Daniel Kaganovich, David W. Taylor, Jack P. K. Bravo, Marianna Teplova, Triana Amen, Eugene V. Koonin, Dinshaw J. Patel, Franklin L. Nobrega

## Abstract

Bacteria and archaea deploy diverse, sophisticated defence systems to counter virus infection, yet many immunity mechanisms remain poorly understood. Here, we characterise the Kiwa defence system as a membrane-associated supercomplex that senses changes in the membrane induced by phage infection and plasmid conjugation. This supercomplex, comprising KwaA tetramers bound to KwaB dimers, as its basic repeating unit, detects structural stress via KwaA, activating KwaB, which binds ejected phage DNA through its DUF4868 domain, stalling phage DNA replication forks and thus disrupting replication and late transcription. We show that phage-encoded DNA mimic protein Gam, which inhibits RecBCD, also targets Kiwa through KwaB recognition. However, Gam binding to one defence system precludes its inhibition of the other. These findings reveal a distinct mechanism of bacterial immune coordination, where sensing of membrane disruptions and inhibitor partitioning enhance protection against phages and plasmids.

## Introduction

Bacteria and archaea have evolved an enormously diverse repertoire of mechanisms to defend against virus predation^1–3^. Among these defence mechanisms, detailed analysis of widespread defence systems, such as CRISPR-Cas and restriction modification not only illuminated fundamental biological processes but allowed adaption of these systems for transformative applications in biotechnology, such as gene editing, expression regulation, and RNA knockdown exploiting CRISPR effectors Cas9, Cas12, and Cas13^4^.

However, the molecular mechanisms of many prokaryotic defence systems, particularly those discovered recently^5–13^, remain poorly (if at all) understood. This knowledge gap is especially pronounced for membrane-associated systems. The Kiwa defence system is one such example. Kiwa consists of two components: KwaA, an integral membrane protein with four transmembrane helices, and KwaB, which contains a DUF4868 domain with an unknown function. Studies on the Kiwa system from *Escherichia coli* O55:H7 RM12579 have demonstrated its ability to protect against phages such as Lambda and SECphi18^6^. Notably, Kiwa is among the 20 most abundant defence systems across sequenced bacterial genomes^1^, highlighting its widespread utility and hence evolutionary success. However, the molecular mechanisms through which Kiwa detects phage infection and provides protection remain elusive.

In this work, we provide mechanistic insights into the Kiwa defence system, revealing that it operates as a transmembrane supercomplex comprising KwaA tetramer bound to KwaB dimers, as its basic repeating unit. This supercomplex detects membrane disruption caused by phage infection and plasmid conjugation, specifically during or prior to phage DNA ejection. Upon activation, KwaB binds phage DNA via its DUF4868 domain, stalling phage replication forks and disrupting phage replication and late transcription. Although DNA binding by KwaB is not sequence-specific *in vitro*, its membrane localisation through KwaA enables a proximity-driven mechanism that minimises off-target effects and prevents host self-damage. Additionally, we demonstrate that Kiwa and RecBCD cooperate to counteract phage-encoded inhibitors, such as the DNA mimic protein Gam, which can only inhibit one of these systems at a time. This interplay underscores the versatility and resilience of bacterial immune networks, offering a perspective on how bacteria thwart phage-encoded inhibitors while safeguarding their cellular functions.

## Results

### Kiwa provides broad protection against phages and plasmids

To investigate the diversity and abundance of Kiwa, we first searched for Kiwa components in 223,143 prokaryotic genomes from the precomputed annotation at the PADLOC web server^13,14^, which is based on RefSeq v209. Kiwa was identified in 9,582 genomes, including 1,713 *E. coli* genomes. We then dereplicated KwaA and KwaB protein sets with a 90% sequence identity threshold and reconstructed the phylogenies of each of these proteins. We found that, although deep branches were not fully resolved, both KwaA and KwaB could be subdivided into three distinct clusters (**Figure 1A**), where the previously described Kiwa operon from *E. coli* O55:H7 RM12579 falls into cluster 1 (purple). The taxonomic assignments of the dereplicated dataset suggest that Kiwa is particularly prevalent in species of the orders *Bacillales* and *Enterobacterales* (**Figure 1B**). Notably, the phylogenetic history of Kiwa substantially deviates from that of their host species, indicating a major role of horizontal gene transfer (HGT) in shaping its evolution (**Figure 1A**). Furthermore, the partial incongruency between the phylogenies of KwaA and KwaB suggests that, on some occasions, *kwaA* and *kwaB* genes were transferred independently (**Figure 1B**).

**Figure 1.**
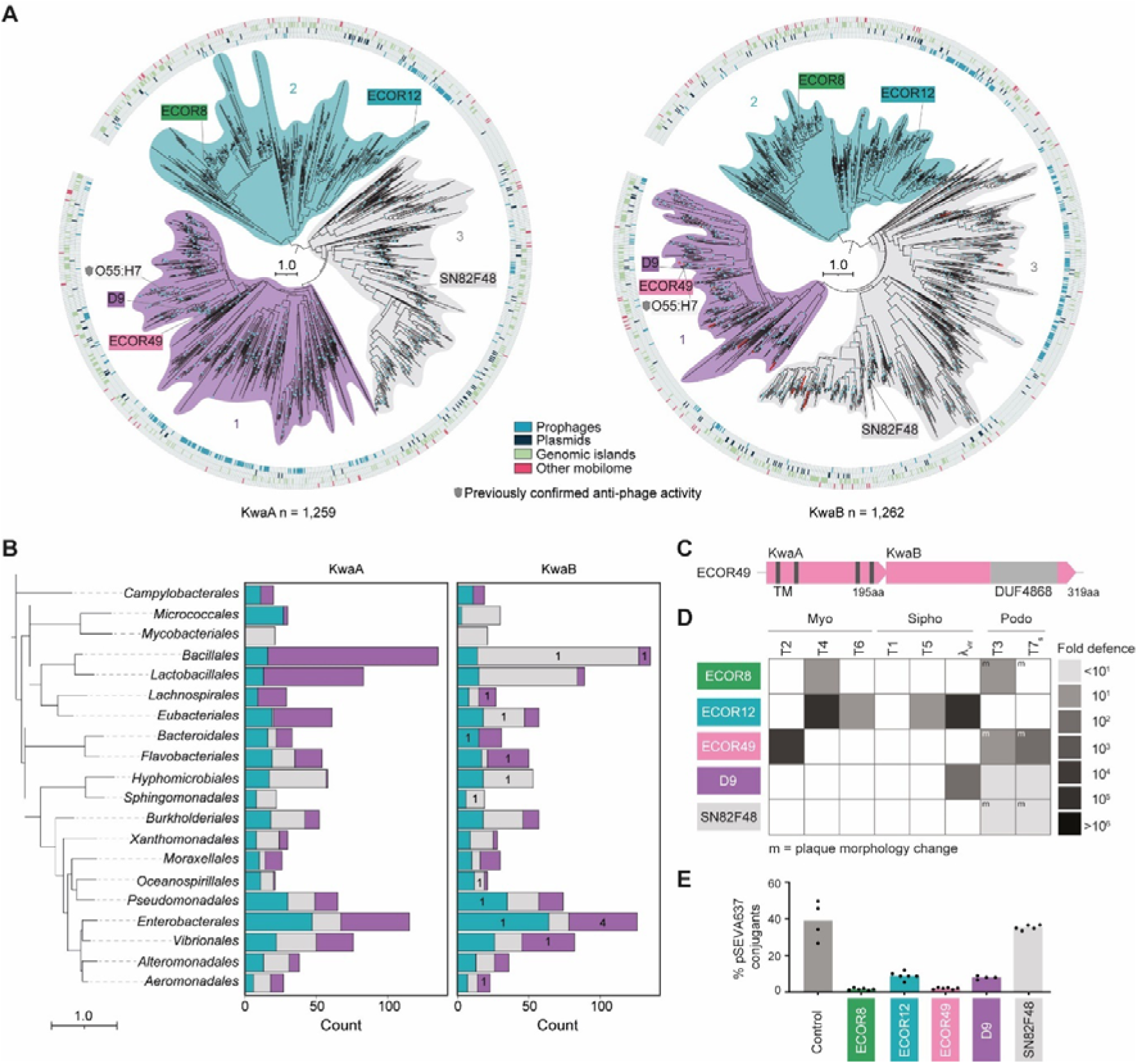
Diversity of Kiwa systems and patterns of defence against phage infection and plasmid conjugation. **(A)** Phylogenetic tree of KwaA (left) and KwaB (right) proteins from the dereplicated dataset. Each cluster is identified by a specific colour shading and numbering. The tree includes annotations for Kiwa operons that were tested in this study, and Kiwa operons that have previously been confirmed to provide anti-phage activity are indicated with a shield symbol. Instances of KwaB split into two genes are annotated in red. The tree scale is in number of substitutions per site and nodes with bootstrap support of at least 80% are highlighted in cyan. Co-localisation of KwaA and KwaB with various classes of MGEs is shown in the out circles. The category “Other mobilome” denotes Kiwa genes that are encoded in close proximity to at least 2 hallmark mobilome genes, as identified by COG annotations^17^, but were not classified as plasmids, prophages, or genomic islands. **(B)** Distribution of KwaA and KwaB proteins across 20 bacterial orders containing the largest number of dereplicated Kiwa components. The bacterial species tree was retrieved from GTDB database^18^, and the scale is in substitutions per site. The barplot was coloured based on phylogenic clusters shown in (A). Numbers in the KwaB barplot indicate instances of KwaB split into two genes. **(C)** Schematic representation of the Kiwa operon obtained from *Escherichia coli* strain ECOR49. TM, transmembrane domain. DUF, domain of unknown function. **(D)** Anti-phage activity of five Kiwa operons shown as fold protection of efficiency of platting assays versus control cells expressing mVenusYFP (YFP). The Kiwa operons were obtained from *E. coli* strains ECOR8, ECOR12, ECOR49, and D9, as well as *Ralstonia mannitolilytica* strain SN82F48. m, plaque morphology change relative to the control strain. **(E)** Anti-conjugation activity of five Kiwa operons shown as percentage of conjugants in relation to total cell count that have uptaken plasmid pSEVA637 from S17-1 donor cells. Bar graphs represent the average of at least four replicates, with individual data points overlaid.

To identify mobile genetic elements (MGE) that might drive HGT of Kiwa operons, we systematically analysed their genomic neighbourhoods. We found that 673 dereplicated *kwaA* (53.5%) and 676 dereplicated *kwaB* (53.6%) genes co-localised with MGEs including 197/198 prophages, 105/110 plasmids, and 304/302 genomic islands for KwaA and KwaB, respectively (**Figure 1A**). In 70 of the 9,797 Kiwa loci, including 19 of the 1,262 in the dereplicated dataset, KwaB is represented by two distinct proteins. The phylogenetic tree topology indicates that KwaB was split into two genes on multiple, independent occasions (**Figure 1A**, right).

Prior studies suggest that diverse variants of the same defence system can have different phage specificities^5,15,16^. To determine if the diversity of Kiwa impacts defence specificity, we cloned five Kiwa systems from different phylogenetic clusters into *E. coli* BL21-AI, a strain naturally lacking Kiwa (**Figure 1C**, **Figure S1A**). When we exposed these strains to a panel of eight phages, all Kiwa systems demonstrated anti-phage activity, each with unique phage specificity and varying levels of protection (**Figure 1D**, **Figure S1B**). We also observed that Kiwa reduced plasmid conjugation efficiency more than twofold, with the exception of Kiwa from *Ralstonia mannitolilytica* strain SN82F48 (**Figure 1E**).

Overall, these findings indicate that Kiwa broadly, albeit with varying specificity, defends bacteria against both phage infections and plasmid conjugation.

### KwaA forms a structurally variable homotetramer

To gain insight into the potential functions of Kiwa components, we first characterised the structure of the transmembrane protein KwaA. Gel filtration indicated that KwaA form an oligomer (**Figure S2A**). Subsequent investigation using cryogenic electron microscopy (cryo-EM) in Lauryl Maltose Neopentyl Glycol (LMNG) detergent revealed two distinct structural states for KwaA, with overall resolutions of 3.7 Å and 4.2 Å, respectively (**Figures S2B,C**; cryo-EM statistics in **Table S1**). These maps allowed us to build a model of the full-length KwaA protein. Side chains were well-defined in the higher resolution 3.7 Å map but less so in the 4.2 Å map (**Figure S2D**).

In both structural states, KwaA forms a homotetramer with an hourglass-like shape (**Figures 2A,B**), which shares similarities with the chloride ion channel TMEM16A. In the absence of calcium ions, TMEM16A homodimers adopt an hourglass-shaped configuration, forming a closed channel. Calcium ion binding near the channel induces a conformational change that opens the channel, allowing negatively charged chloride ions to pass through the membrane^19^. This observation suggests that Kiwa, like TMEM16A, could function as a channel that opens in response to a specific trigger.

**Figure 2.**
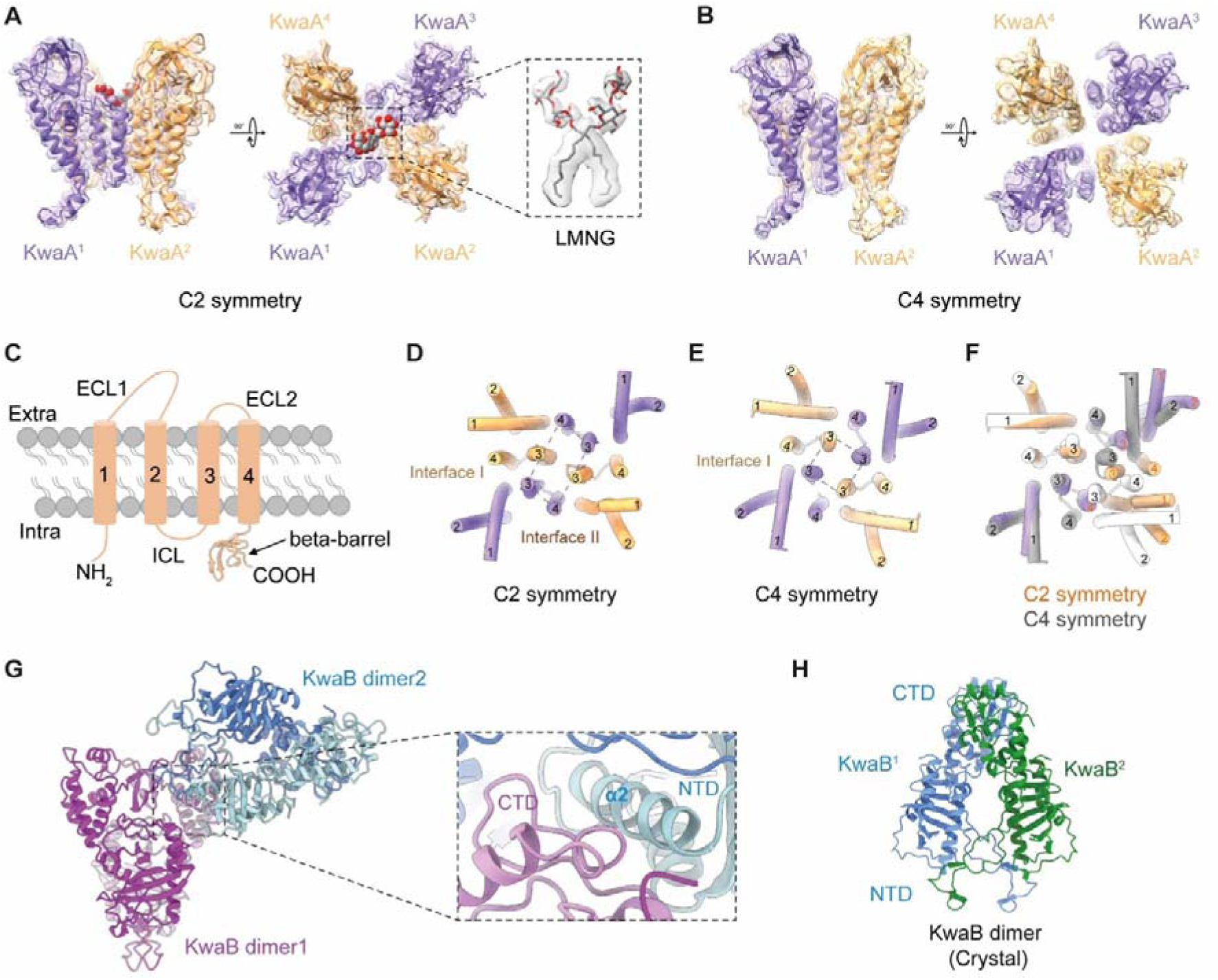
Structures of KwaA and KwaB. **(A)** Two views of the cryo-EM structure of the KwaA tetramer (monomers in purple and orange) adopting C2 symmetry, with an LMNG (boxed segment) inserted into the central channel. **(B)** Two views of the cryo-EM structure of the KwaA tetramer (monomers in purple and orange) adopting C4 symmetry. **(C)** Schematic alignment of transmembrane helices 1 to 4, extracellular loops (ECL1, ECL2), intracellular loop (ICL), and intracellular beta-barrel of a KwaA monomer. **(D)** Helical segments lining the central channel in KwaA tetramer with C2 symmetry, with labelling of interfaces I and II. **(E)** Helical segments lining the central channel in KwaA tetramer with C4 symmetry, with labelling of interface I. **(F)** Structural superposition of helical segments of KwaA tetramers with C2 and C4 symmetry. **(G)** The crystal structure of KwaB. KwaB adopts a dimer of dimers topology with individual dimers coloured in shades of magenta and blue. The expanded boxed segment shows the interface between the two dimers primarily mediated by the C-terminal domain (CTD) of one protomer (in magenta) and the N-terminal α2 helix of another monomer (in blue). **(H)** Ribbon representation of the KwaB dimer (monomers in blue and green) observed in the crystal.

Each KwaA protomer contains four transmembrane helices (TM1-4), two extracellular loops (ECL1-2), one intracellular loop (ICL1), and the intracellular C-terminus, with ICL1 and the C-terminus forming a beta-barrel structure consisting of six beta strands (**Figure 2C**). The two structural states of KwaA differ in their symmetry: the 3.7-Å state exhibits C2 symmetry (**Figure 2A**), whereas the 4.2-Å state has C4 symmetry (**Figure 2B**). In the C2 symmetric structure, the four KwaA protomers adopt two distinct conformations. One conformation resembles the protomers in the C4 symmetric structure (**Figure S2E**, right panel), whereas the other conformation shows notable differences, particularly in the ECL2 and the TM4 regions. In this distinct conformation, TM4 shifts closer to TM3, with a rotation of 30 Å (**Figure S2E**, left panel). This structural rearrangement widens the central pore and repositions the surrounding helices. In the C2-symmetric structure (**Figure 2A**), the pore is formed by TM3 and TM4 from two protomers, along with TM3 from the other two protomers (**Figure 2D**), whereas in the C4 symmetric structure (**Figure 2B**), the pore is exclusively formed by TM3 from all protomers (**Figure 2E**).

Structural alignment of the two states reveals a common interface, designated interface I, which involves interactions between TM1 and TM3 of one protomer and TM3 and TM4 of an adjacent protomer, stabilised by hydrophobic interactions and π-π stacking (**Figure 2F, Figure S2F**). The C2 symmetric structure features an additional unique interface, referred to as interface II (**Figure 2D**), where TM4 from one protomer interacts extensively with TM1 and TM3 of another protomer (**Figure S2F**). This interface exhibits similar hydrophobic and π-π interactions as observed in interface I, emphasizing a conserved assembly mechanism of the KwaA tetramer. The reorganization of ECL2 in the C2-symmetric structure enables connection between two diagonal protomers.

In the expanded view of the central channel in the C2-symmetric structure (**Figure S2G**), we observed a LMNG detergent molecule within the central pore (**Figure 2A**). The two hydrophilic maltoside groups extend outward from the cell membrane, making no distinct interactions with the surrounding KwaA residues, whereas the two hydrophobic chains are deeply embedded into the pore, interacting with nearby hydrophobic KwaA residues (**Figure S2G**).

In summary, the transmembrane KwaA proteins of Kiwa form a homotetramer with two distinct symmetries.

### KwaB forms a dimer of dimers

We next determined the structure of KwaB. The crystal structure at 3.9 Å resolution revealed that KwaB adopts a dimer of dimers topology (**Figure 2G**, **Table S2**). In this configuration, extensive interactions occur between the C-terminal region of one monomer from one dimer (shown in magenta) and the α2 helix of one monomer from another dimer (shown in blue) (**Figure 2G**, expanded view).

KwaB consists of two distinct structural domains: the N-terminal region (residues 1-185), which forms a Rossmann fold-like structure, and the C-terminal region (residues 186-315) that contains a functionally uncharacterised DUF4868 domain (**Figure 2G**, **Figure S3A**). The Rossmann fold-like structure is characterised by a repeating pattern of β-strands alternating with α-helices, forming a three-layered sandwich with a central parallel β-sheet in a βαβ motif. This fold is commonly found in proteins that bind nucleotides including NAD, FAD, oligoA and others, and is one of the most prevalent protein folds, capable of accommodating diverse ligands^20,21^. Both domains contribute to the formation of the KwaB dimer (**Figure 2H**, interactions shown in detail in **Figure S3A,B**).

Our phylogenetic analysis showed that in some lineages, KwaB is split into two distinct genes (**Figure S1A**). To investigate how this affects the functional architecture of KwaB, we predicted the structures of the split proteins using AlphaFold2 and compared them to the resolved structure of single-protein KwaB. This comparison revealed that the split KwaB proteins are closely similar to the two domains of the single-protein KwaB (Rossmann-like fold and DUF4868) (**Figure S3C-E**). Additionally, AlphaFold2 modelling of multimeric structures showed that the tetrameric assembly of split KwaB closely resembles the dimeric architecture of single-protein KwaB (**Figure S3F-H)**. This conservation suggests that the split KwaB proteins evolved via multiple, independent fissions of the ancestral single-protein counterparts, while retaining their domain folds and functional equivalence.

In summary, KwaB forms a robust dimer of dimers structure with multiple stabilising interactions across both the N- and C-terminal regions.

### KwaA and KwaB form a fence-like transmembrane supercomplex

To investigate the roles of KwaA and KwaB and their possible interaction in anti-phage defence by Kiwa, we tested the effect of single-gene deletions. Deletion of either gene resulted in loss of defence, indicating that both are essential for protection (**Figure 3A**). Additionally, truncating the TM domains in KwaA abolished defence, highlighting the importance of membrane localization of KwaA (**Figure 3A**).

**Figure 3.**
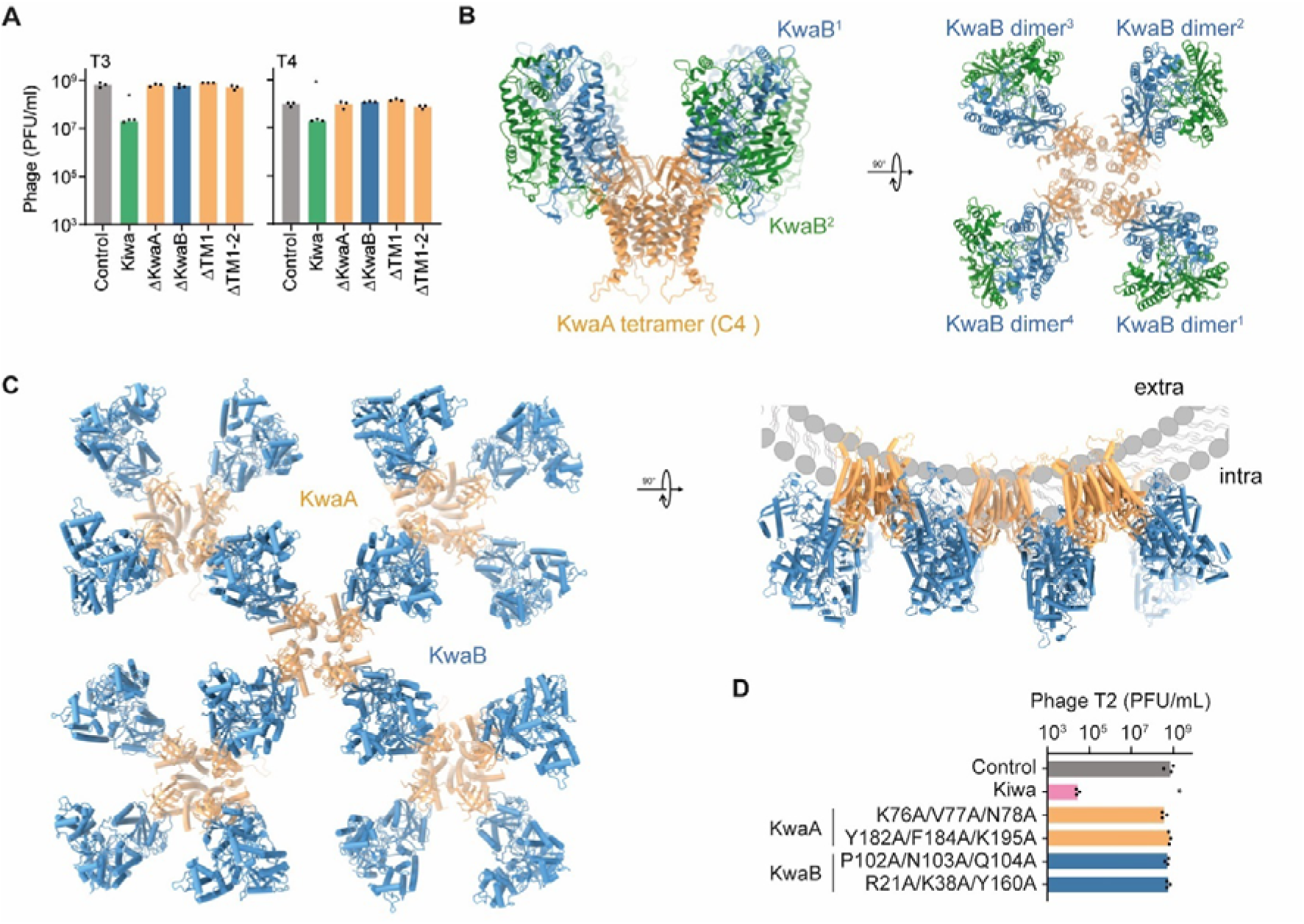
KwaA and KwaB form a supercomplex. **(A)** Impact of deletions and truncations on Kiwa ECOR8 defence against phages T3 and T4 infection. Data represents phage PFU/mL on tested strains, with control expressing YFP. Asterisks indicate a statistically significant increase in PFU/ml compared to Kiwa (Two-way ANOVA, p <0.05). The bars represent the average of three biological replicates with individual data points overlaid. **(B)** Two views of the higher molecular weight structure of the KwaAB complex showing four KwaB dimers (monomers in blue and green) bound to a KwaA tetramer (in orange). **(C)** Structural modelling of a fence-like KwaAB supercomplex on a curved cell membrane with KwaB dimers (in blue) bound to KwaA tetramers (in orange). **(D)** Impact of point mutations on KwaA and KwaB ECOR49 defence against phage T2 infection. Data represents phage PFU/mL on tested strains, with control expressing YFP. Asterisks indicate a statistically significant decrease in PFU/ml compared to control (Two-way ANOVA, p <0.05). The bars represent the average of three biological replicates with individual data points overlaid.

To test complex formation between KwaA and KwaB, we co-expressed both proteins and detected low- and high-molecular weight complexes in sizing column fractions (**Figure S4A**), indicating that KwaA and KwaB interact to form a complex. Using cryo-EM, we visualised the high molecular weight KwaAB complex (**Figure S4B**) and found that it adopts a fence-like, higher-order supercomplex structure (**Figure S4C**). Following multiple classification rounds and symmetry analysis, we resolved the minimal structural unit of KwaAB supercomplex at a 3.7 Å resolution (**Figures S4C,D**; cryo-EM statistics in **Table S1**). This unit features a KwaB homodimer (in green and blue) associated with a tetrameric KwaA (in orange), where each KwaA tetramer binds four KwaB dimers (**Figure 3B, S4E)**. Our structural analysis revealed that the homodimeric structure of KwaB in the KwaAB complex closely resembles its standalone crystal structure (RMSD = 0.63 Å, **Figure S4F**). The KwaA monomers in the KwaAB complex all adopt a similar conformation to their C4 symmetric structure (**Figures 3B, S4C,** right panels), except in regions interacting with KwaB, where localised rearrangements occur (RMSD = 1.24 Å; **Figure S4G**).

In the KwaAB supercomplex, the dimer of dimers configuration of KwaB was not observed. Instead, each KwaB dimer was found to interact with two KwaA tetramers (**Figure 3C)**. Each KwaA monomer engages both protomers of a KwaB dimer through two main interfaces. In the smaller interface (blowup box in **Figure S4H**), the flexible DE loop (residues 134-174, between β-sheets D and E) of KwaB interacts with TM2 of KwaA, where Y157 of the DE loop forms a hydrogen bond with K75 of TM2 (**Figure S4H,I**). At the lateral of the C-terminal beta-barrel, α helix 2 (α2) of KwaB lies adjacent to ICL1 of KwaA. Here, Q79 of KwaA potentially forms a hydrogen bond with N42 of KwaB, while N77 interacts with the main chain of α2, further stabilizing the interface. Triple mutations in KwaA residues involved in this interaction (K75/V76/N77, K76A/V77A/N78A in ECOR49 (71.6% similarity to KwaA from O55:H7)) abolished defence, likely by disrupting the proper KwaAB interactions (**Figure 3D**).

In the larger interface (blowup box in **Figure S4I**), extensive contacts form between the KwaB monomer and the C-terminal beta-barrel region of KwaA, whereas a loop near β-sheet C (residues I96-P108) of KwaB is positioned above the beta-barrel channel. In this area, P100 from KwaB is the deepest-embedded residue, influencing the direction of the loop. This residue is surrounded by H156, Y181 and F183 from KwaA, forming intricate π-π interactions. H156 engages in a cation-π interaction with R21 from beta-sheet A of KwaB, the latter likely forming a hydrogen bond with Y181 and a salt bridge with D80 of KwaA. Additionally, two charged residues, N101 and Q103, within the same loop also contribute polar interactions, N101 with T162 and Q103 with the main chain of KwaA. Mutations in critical KwaA (C-terminal beta-barrel; Y181/F183/K194, Y182A/F184A/K195A in ECOR49) and KwaB (R21/R35/Y157, R21A/K38A/Y160A in ECOR49; P100/N101/Q103, P102A/N103A/Q104A in ECOR49) residues within this larger interface also eliminated defence, demonstrating the key role of the interface (**Figure 3D**).

The KwaB homodimer serves as the structural core for assembling the KwaAB supercomplex (**Figure 3C**, left panel). Aligning the basic units through KwaB dimers surprisingly revealed that the KwaAB supercomplex adopts a bent conformation in its transmembrane region, opposite to the curvature of the native cell membrane (**Figure 3C**, right panel). This organization explains why the supercomplex maintains a C4 symmetry which, unlike a C2 symmetry, is conducive to forming the higher-order complexes required for the assembly of the membrane-spanning KwaAB structures.

An additional band coeluting with the high-molecular weight KwaAB complex on the gel (**Figure S4A**) corresponded to the heptameric, closed-state K^+^ sensitive mechanosensitive channel MscK^22^ (**Figure S4J**). However, MscK was not required for Kiwa-mediated defence against phage infection, as demonstrated by unchanged activity of Kiwa in MscK-deficient cells (**Figure S4K**).

In conclusion, our findings indicate that Kiwa forms a transmembrane supercomplex composed of KwaA tetramer binding of KwaB dimers, as its basic repeating unit, where KwaB homodimers serve as structural anchors for the KwaAB assembly.

### Kiwa localises to phage infection sites and detects phage-induced membrane disturbance

To elucidate the protective mechanism of Kiwa, we investigated whether it induces an abortive infection (whereby the defence system actively causes death of the infected cell to block phage propagation) or mutual destruction (whereby the infected cell dies, blocking phage propagation, not due to the activity of the defence system but as a result of the phage life cycle) phenotype during phage infection^23,24^. We monitored phage and cell concentrations in liquid cultures at high phage pressure (MOI = 10) and found that Kiwa substantially reduced the amount of phage progeny while preserving viability of most cells (**Figure 4A**). In contrast, control cells lacking Kiwa were reduced to below the detection limit of our assay. These findings indicate that Kiwa does not trigger an abortive infection or mutual destruction but rather appears to protect individual cells from phage infection.

**Figure 4.**
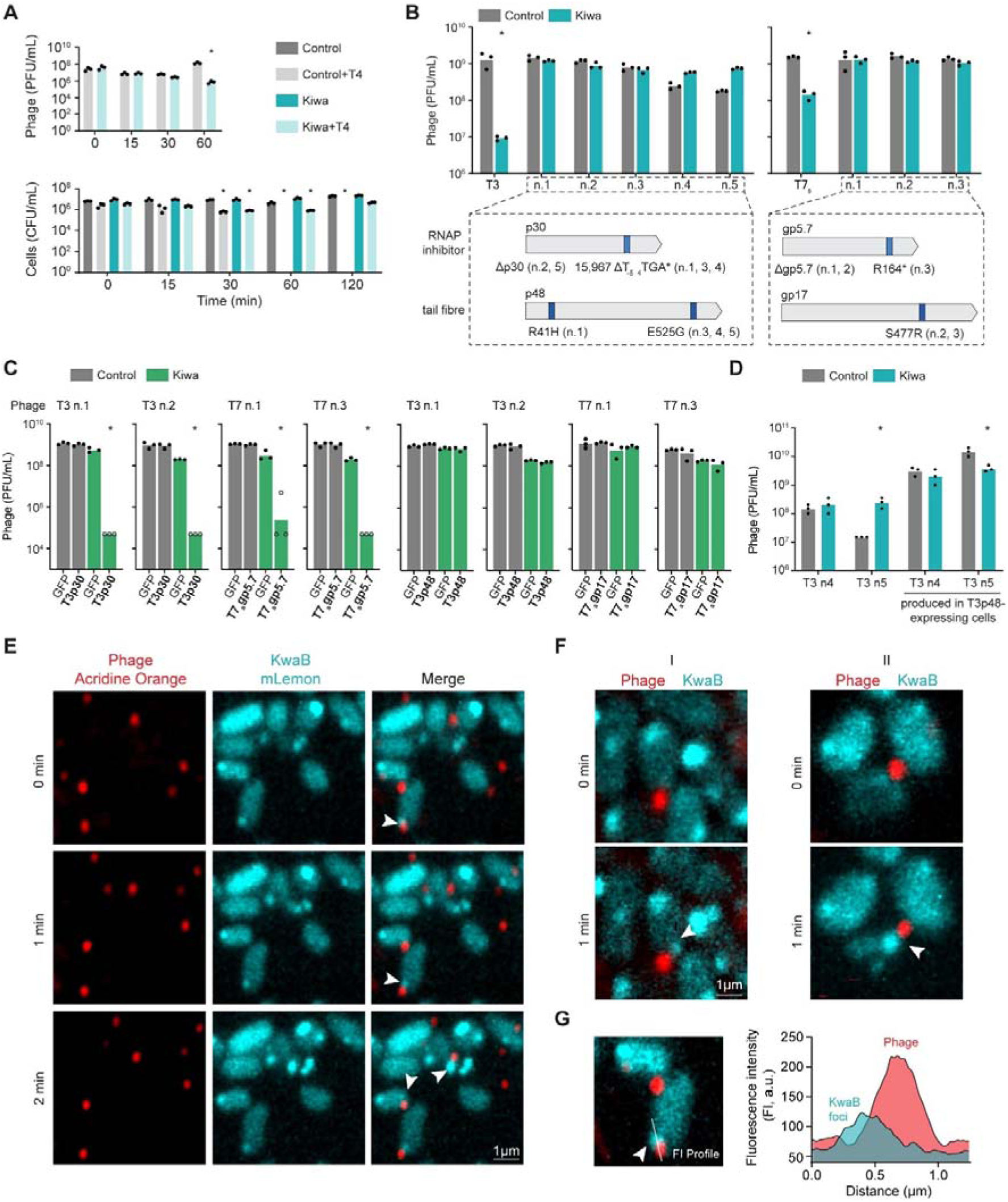
Kiwa localises to phage infection sites before phage DNA ejection occurs. **(A)** Phage and cell titres (PFU/ml and CFU/ml, respectively) measured at different time points post infection of control cells and Kiwa cells with phage at a multiplicity of infection (MOI) of 10. The average of three biological replicates is shown with individual data points overlaid. Asterisks indicate a statistically significant difference in phage or cell titre compared to the titres obtained in control cells (Two-way ANOVA, p <0.05). **(B)** Phage titres (PFU/ml) of wild type (T3, T7s) and mutant phages (n) in control cells expressing YFP and in cells expressing Kiwa. The average of three biological replicates is shown, with individual data points overlaid. Asterisks indicate a statistically significant decrease in phage titre compared to the titre obtained against the control cells (Two-way ANOVA, p <0.05). Mutations found in the escape phages are depicted below the graphics. The full list of mutations and deletions is detailed in Table S3. **(C)** Reconstitution of T3p30 and T7s gp5.7 recovers Kiwa defence activity against phage escape phages, while reconstitution of T3p48 and T7s gp17 did not. Asterisks indicate a statistically significant decrease in phage titres relative to control cells (GFP) (Two-way ANOVA, p <0.05). Bars represent the average of three biological replicates with individual data points overlaid. **(D)** Phage titres of T3 phage escape mutants and T3 escape mutants propagated in cells expressing wild type T3p48 in *trans*, in control and Kiwa cells. Asterisks indicate a statistically significant decrease in phage titres relative to control cells (Two-way ANOVA, p <0.05). Bars represent the average of three biological replicates with individual data points overlaid. **(E)** Confocal microscopy images of Kiwa-expressing cells infected with phage. Fluorescently tagged KwaB (mLemon, depicted in cyan) and SYTOX Orange-stained phage particle DNA (red) are shown. White arrows highlight instances where KwaB foci are proximal to the phage. **(F)** Enlarged view of regions showing KwaB foci forming near phage attachment sites. Observed by confocal microscopy as in (E), these foci appear before phage DNA is (fully) ejected into the cell. **(G)** Fluorescence intensity profiles of KwaB (depicted in blue) and phage (red), demonstrating overlapping signals that indicate spatial proximity.

To further explore the mechanism of Kiwa defence, we isolated phage mutants escaping protection by Kiwa (**Figure 4B**). Sequencing of these escape mutants, from both T3 and T7Select (T7s) phages, revealed mutations in two types of genes: (1) inhibitors of the stationary phase σ subunit (RpoS) of host RNA polymerase (RNAP) (T7s gp5.7 and T3 T3p30)^25,26^ and (2) tail fibre genes (gp17 in T7s and T3p48 in T3) (**Figure 4B**, **Table S3**). To validate the role of these genes in Kiwa activation, we expressed the respective proteins in *trans*. The RpoS inhibitors were found to restore Kiwa protection against escape phages, but the tail fibre genes did not (**Figure 4C**).

We hypothesised that the tail fibre proteins of infecting phage particles might externally activate Kiwa. To test this proposition, we propagated the escape mutant T3 phages in cells expressing wild-type T3p48 *in trans*, enabling some newly produced phage particles to incorporate the wild-type T3p48 protein. Mutants T3 n4 and n5 were chosen for their high efficiency in escaping Kiwa defence (**Figure 4B**). Despite low tail fibre incorporation efficiency, we expected that differences in infectivity could be detected. When these mutant T3 populations, likely containing a small fraction of phages with wild-type T3p48, were used to challenge Kiwa-expressing cells, mutant n5 triggered Kiwa defence. This finding confirmed that tail fibre proteins can activate Kiwa from outside the cell (**Figure 4D**).

Given that the KwaAB transmembrane supercomplex adopts a curvature opposite to the normal membrane curvature (**Figure 3C**) and appears to be activated externally, we hypothesised that Kiwa sensed changes at the membrane level. For well-characterised phages like T4, DNA ejection is known to involve the formation of a membrane-bound channel by the phage tail, causing outward bulging of the cytoplasmic membrane^27^. The large size and mesh-like structure of the KwaAB supercomplex, which spans a broad membrane area, could allow Kiwa to detect membrane curvature changes caused by phage or pilus attachment. If this hypothesis is correct, Kiwa should co-localise with the site of phage infection. To test this, we performed co-localisation assays using a Kiwa system with fluorescently tagged KwaB (mLemon) and phage particles stained with DNA dye SYTOX orange. KwaB foci consistently formed near the site of phage binding, at the cell poles, before DNA ejection occurred, as indicated by the DNA dye remaining within the phage capsid (**Figure 4E-G**, **Figure S5**). Quantification of phage particles on cells in proximity to KwaB foci showed that 26 ± 3% of the phages are proximal to KwaB (defined by the partial overlap of fluorescent intensity profiles). Given the dynamic nature of this process, this percentage aligns with expectations. These results further support the idea that phage binding or the initiation of DNA ejection triggers Kiwa defence.

In summary, these findings indicate that Kiwa defence is initiated at the site of phage attachment, likely triggered by early structural changes such as membrane curvature during phage binding or DNA ejection.

### KwaB inhibits phage late transcription

To elucidate the role of KwaB in defence, we examined the structure of this protein more closely. Structural analysis of KwaB (**Figure 2G,H**) revealed an N-terminal Rossmann fold, a common structural domain that typically binds nucleotide second messengers, triggering a conformational change in the respective proteins that activates an effector domain^28^. Homology-based functional predictions using SPROF-GO^29^ suggested that KwaB binds nucleic acids, which is supported by electrostatic analysis of the KwaB dimer. This analysis revealed a positively-charged central channel between the two KwaB monomers (**Figure 5A**), suggesting that this site might bind negatively charged molecules, such as DNA.

**Figure 5.**
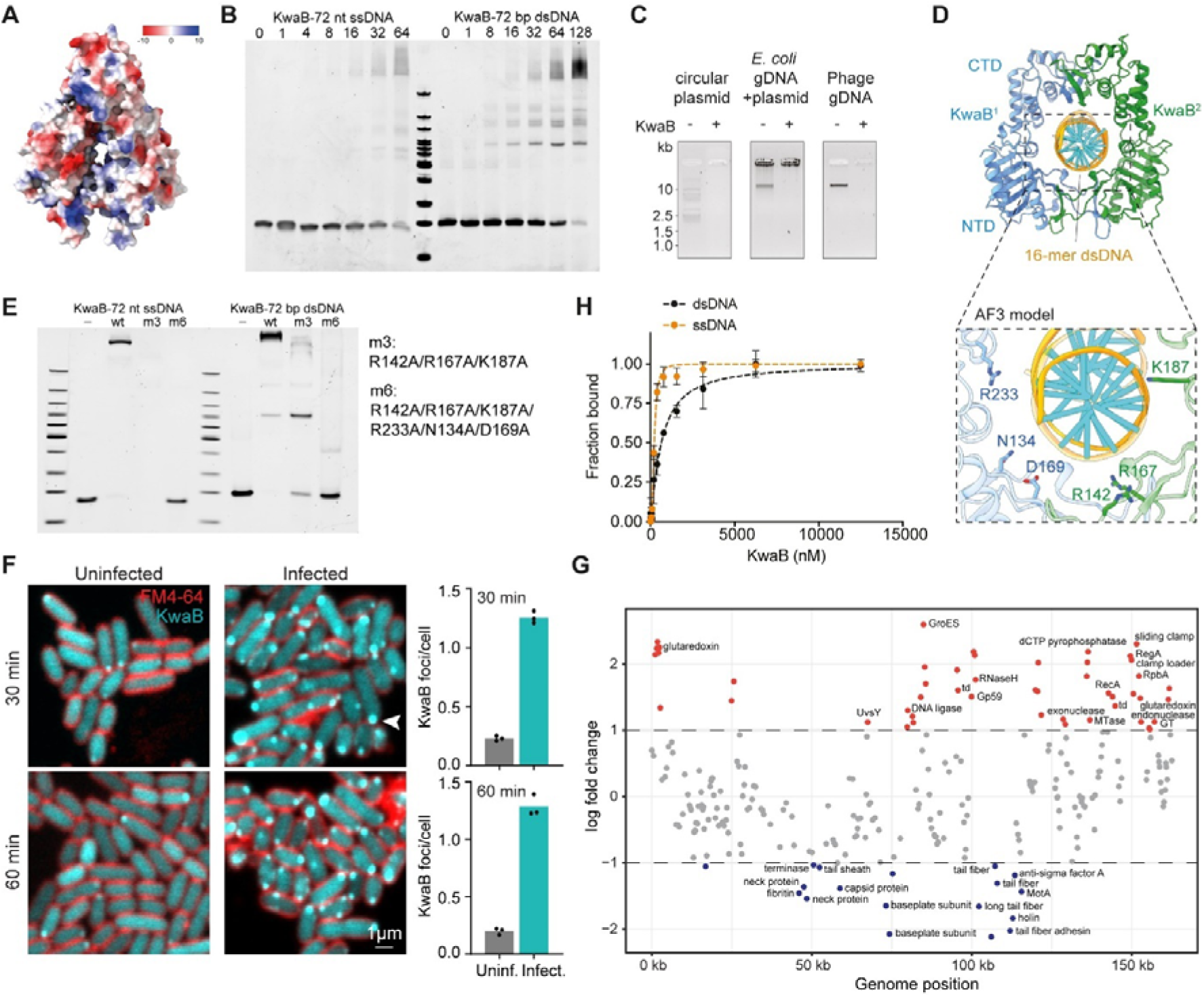
KwaB binds to phage DNA and reduces late gene transcription. **(A)** Electrostatic surface representation of KwaB dimer. **(B)** Electrophoretic mobility shift assays with KwaB and a 72 nucleotide ssDNA and dsDNA target. Increasing concentrations of KwaB (μM, show on top of the gel) were incubated with 10 nM ssDNA or dsDNA. **(C)** Electrophoretic mobility shift assays with KwaB and circular plasmid DNA, genomic bacterial DNA, and genomic phage DNA. 500 ng of KwaB were incubated with 50 ng DNA. **(D)** Alphafold 3 (AF3) model of the KwaB dimer (monomers in blue and green) complexed with a 16-mer dsDNA, highlighting positioning of the dsDNA between two KwaB monomers. Protein-DNA contacts are highlighted in the expanded boxed segment. **(E)** Impact of mutations in residues involved in KwaB-DNA contacts on KwaB binding to dsDNA. Electrophoretic mobility shift assays were performed with mutant versions of KwaB (2.56 μM) with a 72 bp ssDNA or dsDNA target (10 nM), as in (B). **(F)** KwaB forms membrane-localised foci throughout phage infection. On the left, confocal microscopy images on the left show Kiwa cells with mLemon tagged KwaB (depicted in blue) uninfected and infected for 30 and 60 minutes. Membrane is stained with FM4-64 (red). On the right, quantification of KwaB foci per cell. Bars represent the average with individual points overlaid. **(G)** Log fold change in gene transcription in Kiwa-expressing cells compared to control cells infected with phage T2 at 15 min. Genes overtranscribed in Kiwa cells are shown in red and those undertranscribed are shown in green. **(H)** Fluorescence anisotropy binding assays of KwaB to a 20-bp ss- or ds-DNA substrate. Data represents the bound fraction of substrate relative to KwaB (nM). Data shows the average and standard deviation of three independent replicates.

To test this hypothesis, we examined the DNA-binding capacity of KwaB using electrophoretic mobility shift assays (EMSA). Band shifts confirmed that KwaB binds both linear ssDNA and linear dsDNA (**Figure 5B**). Moreover, KwaB also binds circular plasmid dsDNA, bacterial genomic dsDNA (gDNA), and phage gDNA (**Figure 5C**), indicating a broad, non-specific DNA binding affinity *in vitro*. Importantly, we observed no degradation products from either dsDNA or ssDNA targets (**Figure S6A**), indicating that KwaB binds but does not degrade nucleic acid substrates.

For a closer examination of KwaB-DNA interactions, we used AlphaFold3 (AF3) to predict the structure of KwaB dimer bound to dsDNA. As anticipated, the model demonstrated dsDNA threading through the positively charged central channel of the KwaB dimer, where interactions with surrounding residues (R233, N134, D169, R142, R167, K187) stabilise binding (**Figure 5D**). We observed a conformational shift between KwaB monomers when transitioning from the apo state (**Figures 2H, 3B**) to the dsDNA-bound state (**Figures 5D**), reflecting opening of the channel to accommodate the bound dsDNA. The residues predicted to interact with dsDNA are located in the DUF4868 domain of KwaB, whose function remains unclear. Here, we show that mutations of all DNA-contacting residues (m6: R142A/R167A/K187A/R233A/N134A/D169A) severely reduced KwaB binding to ssDNA and dsDNA (**Figure 5E**, **Figure S6B**), indicating a critical role of DUF4868 in DNA binding.

We hypothesised that KwaB captures the ejected phage DNA at the bended membrane, preventing access to the DNA replication machinery. To test this hypothesis, we used confocal microscopy which confirmed that KwaB remains membrane-associated, at the cell poles, via its interaction with KwaA throughout phage infection (**Figure 5F**, **Figure S6C**). As the localised foci formed by fluorescently tagged KwaB in close proximity to the site of phage DNA entry, these findings suggest that KwaB may interact with the ejected phage DNA, interfering with its transcription and/or replication (**Figure 4E-G**).

To explore this further, we performed RNA-seq of T2-infected Kiwa and control cells at 15 minutes post-infection. Of the 255 phage genes, 187 showed unchanged expression levels, whereas 18 late genes were substantially under-expressed (log2-fold change < -1) in Kiwa-containing cells. These under-expressed genes encode components critical for phage morphogenesis (e.g. structural proteins, terminase), indicating specific failure of late transcription (**Figure 5G**, **Table S4**). Conversely, 59 phage genes were over-expressed in Kiwa cells. These included genes involved in DNA replication (e.g., helicase loader, clamp loader), DNA repair and recombination (UvsY, thymidylate synthase, RecA), DNA modification (Dam methyltransferase), and late transcription activation (e.g., GP45.1, RpbA, RegA). For T4-like phages such as T2, late transcription depends on ongoing DNA replication, whereas RNA polymerase transcribes newly replicated DNA, tightly coupling replication to late gene expression^30^. Therefore, late transcription in Kiwa-expressing cells likely fails because KwaB DNA binding stalls replication forks, preventing the synthesis of the DNA templates required for late-stage transcription. The local, transient ssDNA-dsDNA junctions at replication forks may serve as specific interaction targets for KwaB given that this protein shows affinity to both DNA forms, with a preference for ssDNA (**Figure 5B,H**).

To directly test for replication defects, we quantified phage DNA levels in Kiwa and control cells. Phage DNA levels were reduced by approximately 25% in Kiwa-expressing cells compared to controls (**Figure S6D**), consistent with stalled replication. This defect coincides with the overexpression of phage DNA repair and recombination genes, which are essential for reassembling functional replication forks^31^. The upregulation of these pathways suggests that phage genomes experience replication stress, likely caused by KwaB binding to DNA which interferes with fork progression. This replication block explains the failure to produce late transcripts, despite the upregulation of late transcription activators and proteins promoting replication.

In summary, these results demonstrate that KwaB binds ejected phage DNA via the DUF4868 domain and disrupts replication fork progression, stalling phage DNA replication, preventing late transcription, and ultimately blocking phage morphogenesis.

### Kiwa is antagonised by phage DNA mimic protein Gam

Phages often encode proteins that counteract bacterial defence systems, allowing successful phage propagation^32,33^. Among these proteins are DNA mimics that prevent DNA-targeting defence mechanisms from accessing phage genomes^34–36^. Because we observed DNA binding by KwaB, we hypothesised that DNA mimics might interfere with the anti-phage activity of Kiwa.

To investigate this possibility, we tested the effect of two well-known DNA mimic proteins – Ocr from phage T7^36^ and Gam from phage lambda^37^ – on Kiwa defence against phage T4, which lacks these inhibitors. Infection assays demonstrated that Gam inhibited Kiwa, as indicated by an approximately 100-fold decrease in protection (**Figure 6A**). This was further validated by assays using a Lambda phage with an inactive Gam protein^38^, against which Kiwa was approximately 100-fold more effective than against wild-type Lambda (**Figure S7A**).

**Figure 6.**
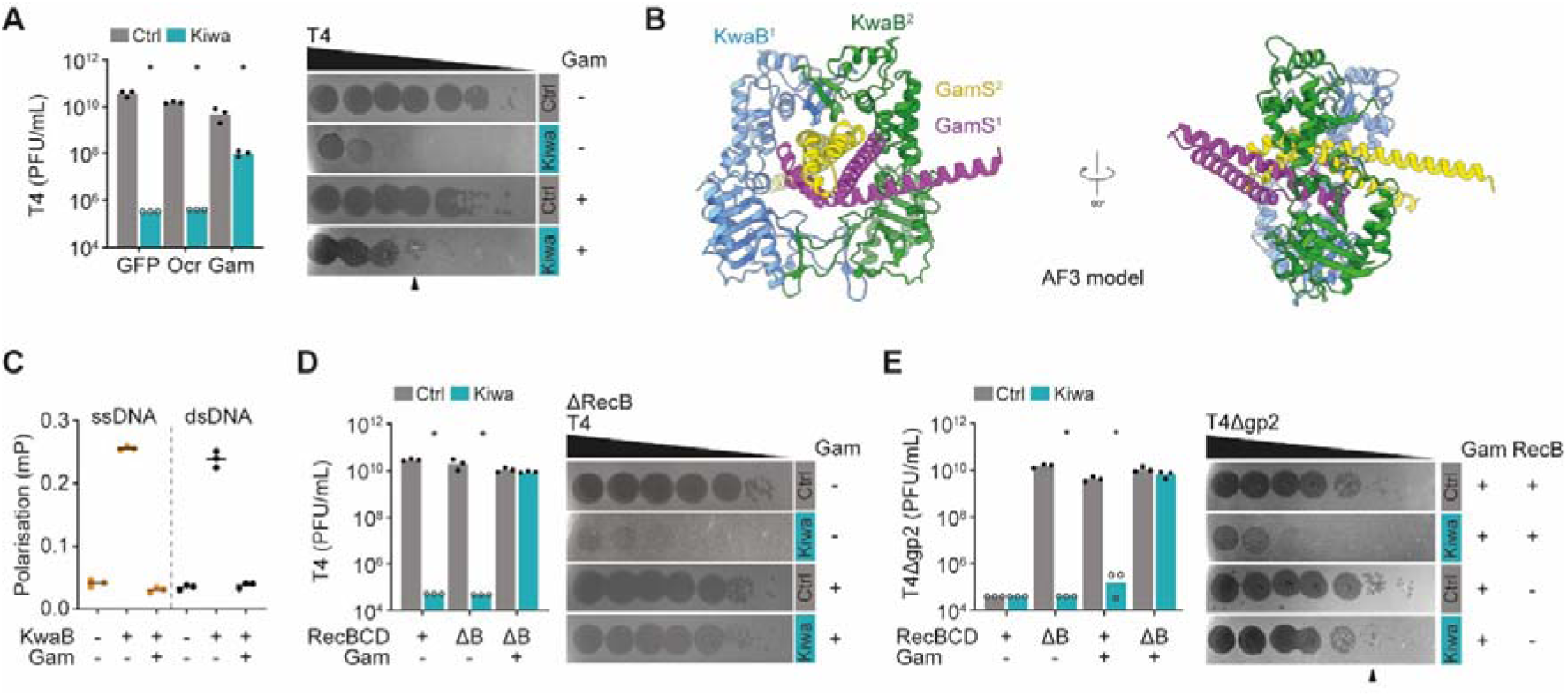
Kiwa is antagonised by phage DNA mimic protein Gam. **(A)** Effect of phage T7 Ocr and Lambda Gam protein expression on Kiwa anti-phage activity against phage T4. Data on the left represents phage PFU/mL on control (YFP (Ctrl)) or Kiwa strains with indicated proteins expressed in *trans*. Asterisks indicate a statistically significant decrease of phage concentration (PFU/mL) relative to control cells expressing the same protein in *trans* (Two-way ANOVA, p <0.05). Bars represent the average of three biological replicates, with individual data points overlaid. Open points indicate instances where it was not possible to determine individual phage plaques, hence a value of 1 was assumed at the corresponding dilution. On the right, individual representative images are shown of spot assays of phage T4 dilutions on Ctrl or Kiwa strains with or without Gam. Indicated is the dilution where individual plaques are observed on Kiwa Gam+, whereas they were not on Kiwa Gam-. **(B)** Two views of AlphaFold3 (AF3) predicted model of the KwaB dimer (monomers in blue and green) complexed with a GamS dimer (in magenta and yellow). **(C)** Fluorescence anisotropy binding of KwaB to a 20 bp ssDNA or dsDNA substate in the presence or absence of Gam. Data represents polarisation of substrate in millipolarisation (mP), showing three representatives with individual points overlaid. Experiments were performed using 1 µM KwaB with 50 nM ss- or dsDNA and incubated with or without 10 µM of Gam protein. **(D)** Effect of RecB subunit deletion and Gam expression on Kiwa anti-phage activity against phage T4. Data on the left represents phage PFU/mL on control or Kiwa cells with intact RecBCD or deleted RecB (ΔB), in the absence and presence of Gam. Asterisks indicate a statistically significant decrease of phage concentration (PFU/mL) relative to control cells on the same host strain (Two-way ANOVA, p <0.05). Bars represent the average of three biological replicates, with individual data points overlaid. Open points indicate instances where it was not possible to determine individual phage plaques, hence a value of 1 was assumed at the corresponding dilution. On the right, individual representative images of spot assays of phage T4 dilutions on ΔRecB Ctrl or Kiwa strains with or without Gam. **(E)** Effect of ΔRecB and Gam expression on Kiwa anti-phage activity against phage T4Δgp2. Data on the left represents phage PFU/mL on control or Kiwa strains with complete RecBCD or RecB deletion (ΔB), in the presence or absence of Gam expression. Asterisks indicate a statistically significant decrease of phage concentration (PFU/mL) relative to control cells on the same host strain (Two-way ANOVA, p <0.05). Bars represent the average of three biological replicates, with individual data points overlaid. Open points indicate instances where it was not possible to determine individual phage plaques, hence a value of 1 was assumed at the corresponding dilution. On the right, individual representative images of spot assays of phage T4Δgp2 on Ctrl or Kiwa strains with or without Gam, and with or without RecB. Indicated is the dilution where individual plaques are observed on Kiwa Gam+ RecB-but not on Kiwa Gam+ RecB+.

To explore the interaction between Gam and KwaB, we performed BACTH (bacterial adenylate cyclase-based two-hybrid) assays with KwaB-T18 and T25-Gam (**Figure S7B**) and co-purification experiments (**Figure S7C**), both of which suggested potential interaction. Although we could not resolve the structure of Gam bound to the KwaB dimer due to sample aggregation and precipitation, AF3 modelling predicted a binding mode for the KwaB-Gam complex resembling the interaction of KwaB with DNA (**Figure 5D**). The model shows a pair of Gam homodimers (in yellow and magenta) inserting into the central positively charged channel of KwaB (**Figure 6B**). As with dsDNA binding (**Figure 5D**), AF3 models predict that Gam binding induces a conformational shift in KwaB, changing its shape from a V- to U-shaped topology (**Figure 2H**, **Figure 6B**).

To determine if Gam competes with DNA for KwaB binding, we conducted competition assays using KwaB, ss- or dsDNA, and Gam. For both DNA substrates, Gam binding to KwaB abolished KwaB binding to DNA, as indicated by decreased polarisation of the DNA target upon Gam addition (**Figure 6C**).

Together, these findings indicate that the Gam protein inhibits Kiwa defence by binding to KwaB and blocking its access to DNA.

### Co-occurrence of Kiwa and RecBCD enables phage defence in the presence of Gam

The Gam protein of phage Lambda inhibits the bacterial RecBCD complex by directly binding to the RecB subunit^39^. RecBCD plays a critical role in bacterial recombination and immunity, degrading phage DNA^40,41^, and supporting other defence systems such as CRISPR-Cas^42^ and prokaryotic Argonaute^43^. Given that RecBCD is present in cells used to assay the inhibition of Kiwa by Gam, we hypothesized that Kiwa was only partially inhibited due to competition between Kiwa and RecBCD for Gam binding. To investigate this possibility, we tested the effect of Gam on Kiwa on a ΔRecB strain, effectively removing one of Gam’s targets.

Whereas Gam only partially inhibited Kiwa in RecB+ cells, it completely suppressed Kiwa in ΔRecB cells (**Figure 6D**), indicating that the inhibitory effect of Gam on Kiwa is enhanced when a competing target is absent. This observation was reinforced by infection assays using a wild type Lambda and a variant with an inactive Gam^38^, where Kiwa activity remained robust in the absence of RecB against the Gam-inactive variant but was reduced against the Gam-containing phage (**Figure S7A**).

To further assess the contribution of RecBCD to Kiwa defence, we used a T4 mutant lacking the DNA-end protecting protein gp2, which prevents RecBCD-mediated DNA degradation^44^. Infection assays showed that RecBCD targets T4Δgp2 as efficiently as Kiwa does (**Figure 6E**). Furthermore, although Gam inhibited RecBCD in control cells, Kiwa (+RecBCD) retained activity despite the presence of Gam (**Figure 6E**). By contrast, in the absence of RecB, Gam completely abolished Kiwa protection against T4Δgp2 (**Figure 6E**). Importantly, Gam inhibition of RecBCD and Kiwa is structurally exclusive so that a single Gam dimer cannot inhibit both systems simultaneously (**Figure S7D**).

Overall, our results show that, although RecBCD and Kiwa are each individually susceptible to Gam inhibition, their coexistence confers resilience, preventing Gam from fully suppressing their functions.

## Discussion

In this study, we describe the structure and detailed mechanism of the Kiwa defence system, a membrane-embedded KwaAB supercomplex found in a broad range of bacterial genomes. Our data indicate that Kiwa appears to operate by sensing changes in the membrane caused by phage attachment or plasmid conjugation. The detection of membrane change, mediated by the transmembrane protein KwaA, triggers KwaB, which binds ejected phage DNA, apparently disrupting phage DNA replication and consequently late transcription.

Sensing of membrane changes by KwaA is central to Kiwa activation, reflecting a broader role of membrane sensing in both prokaryotic and eukaryotic cellular responses to structural stress^45^. The DNA-binding of KwaB, mediated by its DUF4868 domain, is not sequence-specific *in vitro*, yet its localisation to the membrane via KwaA enables a targeted, versatile anti-phage response that prevents phage DNA replication and late gene transcription. This proximity-driven defence strategy seems to limit off-target effects, sparing the host from potential self-damage. In addition, DNA-binding by KwaB might be specifically activated by conformational changes in the KwaAB complex, triggered by membrane distortion caused by phage or plasmid, which would further safeguard the host against self-targeting.

A key finding of our study is the susceptibility of Kiwa to phage-encoded DNA mimicry proteins, such as Gam from phage lambda^46^, which inhibits Kiwa by binding to KwaB. We observed that, whereas both Kiwa and RecBCD defences are individually vulnerable to Gam inhibition, their co-existence in *E. coli* provides a layer of resilience. Specifically, Gam only partially inhibits Kiwa in the presence of RecBCD, suggesting that Gam proteins are functionally partitioned between these two defence targets. In RecBCD-deficient cells, however, Gam fully suppresses Kiwa, indicating that these overlapping defences help to prevent phage evasion of immunity by conferring redundancy and hence robustness on the immune repertoire of the bacterial cell.

This redundancy underscores the adaptive advantage of the presence of multiple defence systems in bacterial cells. Overlapping defences protect against single-point failures in the presence of phage inhibitors, preventing a single inhibitor from entirely bypassing host immunity. While harbouring multipurpose inhibitors, such as DNA mimics^35,47–49^, might benefit phages by conserving genomic resources, the simultaneous presence of targeted defence systems, such as RecBCD and Kiwa, in the host cell maintains an effective barrier against such inhibitors and thus against phage reproduction.

Altogether, our findings reveal how bacteria have evolved to detect and respond to changes in the membrane caused by phage infection or plasmid conjugation. Furthermore, our findings support the view that bacterial defence systems operate as interconnected, multilayered networks rather than isolated modules, leveraging redundancy and versatility to maintain defence efficacy amidst rapidly evolving phage threats.

## Acknowledgements

We thank Rotem Sorek from the Weizmann Institute of Science for kindly providing a Lambda phage Gam mutant, and Ian Molineux from the University of Texas for kindly providing phage T4Δgp2. We also thank YouYu from Zhejiang University-University of Edinburgh Institute and J. De La Cruz (MSK) for assistance with cryo-EM data collection, and Lyuqin Zheng (MSK) for discussions on structural analysis. We thank the Imaging and Microscopy centre (IMC) facility at the University of Southampton. This work was supported by grant RGS\R2\222312 from the Royal Society to F.L.N., Welch Foundation grant F-1938 to D.W.T., and the National Institutes of Health R35GM138348 to D.W.T. T.A. is supported by Wessex Medical Research Innovation Grant (AE06). D.J.P. is supported by NIH grant GM145888, the Maloris Foundation and Memorial Sloan-Kettering Core grant (P30-CA008748). In addition to access of the cryo-EM resources at MSKCC, some of this work was performed at the National Center for CryoEM Access and Training (NCCAT) and the Simons Electron Microscopy Center located at the New York Structural Biology Center, supported by the NIH Common Fund Transformative High Resolution Cryo-Electron Microscopy program (U24 GM129539,) and by grants from the Simons Foundation (SF349247) and NY State Assembly. This research used NSLS-II MX x-ray User Resources (FMX) of the National Synchrotron Light Source II, a U.S. Department of Energy (DOE) Office of Science User Facility operated for the DOE Office of Science by Brookhaven National Laboratory under Contract No. DE-SC0012704. The Center for BioMolecular Structure (CBMS) is primarily supported by the National Institutes of Health, National Institute of General Medical Sciences (NIGMS) through a Center Core P30 Grant (P30GM133893), and by the DOE Office of Biological and Environmental Research (KP1605010). R.K. and E.V.K. are supported by the Intramural Research Program of the National Institutes of Health of the USA (National Library of Medicine).

## Author contributions

Conceptualization, F.L.N.; Methodology; Z.Z., T.C.T., Y.W., R.K., D.K., E.V.K., D.P., and F.L.N.; Formal Analysis, T.C.T., T.A., R.K., F.L.N., J.P.K.B, and D.W.T.; Investigation, Z.Z., T.C.T., Y.W., J.C.R., M.T., T.A., A.N., R.M., and J.P.K.B; Visualization, Z.Z., T.C.T., T.A., Y.W., R.K., and T.A.; Data Curation, T.C.T., Y.W., F.L.N.; Writing – Original Draft, Z.Z., T.C.T., Y.W., R.K., E.V.K., D.P., and F.L.N.; Writing – Review & Editing, D.P., E.V.K., and F.L.N.; Resources, D.K., D.W.T., D.P., E.V.K., and F.L.N.; Funding Acquisition, F.L.N..

## Methods

### Data availability

All unique bacterial strains, phages, and plasmids generated in this study are available from the lead contact without restriction. Raw data have been deposited at Figshare, Protein Data Base (PDB) and the Electron Microscopy Data Bank (EMDB) and are publicly available as of the date of publication.

### Bacterial strains and phages

*E. coli* Dh5α, BL21-AI, BL21-AIΔRecB, BL21-AIΔRecC, BL21-AIΔRecD^42^, KEIO BW25113 and JW0454^50^ were grown at 37 °C in Lysogeny Broth (LB) for liquid cultures or LB agar (LBA, 1.5 % (w/v) agar) for solid cultures. Whenever applicable, LB was supplemented with chloramphenicol (25 µg/mL), spectinomycin (50 µg/mL), Isopropyl ß-D-1-thiogalactopyranoside (IPTG, 1 mM) or L-Arabinose (0.2 %), to ensure maintenance of plasmids or induce protein expression. Phage infection was performed in LB at 37 °C. Phages were propagated by infecting *E. coli* BL21-AI grown to an optical density at 600 nm (OD_600_) of 0.2-0.4 with 5-10 µL of a phage lysate. Infected cultures were grown for at least 4 hours or until culture collapse. The culture was then centrifuged at 11,000 × *g*, 4 °C for 15 min to remove cell debris, and the supernatant was filter sterilized through a 0.22 µm filter. Phage lambda with a frameshift mutation in Gam^38^ was produced in BL21-AI ΔRecB. All strains and phages used in this study are listed in **Table S5**.

### Plasmid and strain construction

Kiwa systems from *E. coli* ECOR8, ECOR12, ECOR49, and D9 were amplified by Q5 polymerase (New England Biolabs) using primers indicated in **Table S6**. Kiwa from *Ralstonia mannitolilytica* SN82F48 was ordered synthetically as a gBlock (Integrated DNA Technologies) (**Table S6**). Kiwa systems under with native promoters were amplified from the respective host strain using primers listed in **Table S6**, in reactions that added regions of homology for cloning into pUOS016 using NEBuilder Hifi DNA Assembly Mastermix. When appropriate, deletions, mutations and truncations of Kiwa operons were engineered by around-the-horn PCR with primers listed in **Table S6**. For protein purification, His-tagged KwaB of ECOR8 or ECOR12 Kiwa was constructed by around the horn PCR on pUOS017 or pUOS018 respectively, introducing a 9xHis tag at the N-terminus of KwaB with primers listed in **Table S6**. Phage proteins gp5.7/T3p30 and gp17/T3p48 were amplified from the genome of T7s or T3 respectively using the primers indicated in **Table S6**. These proteins were cloned into pCDF1-b (Novagen) under the control of a T7 promoter. Lambda GamL was amplified from Lambda genomic DNA and cloned into pCDF1-b by Gibson assembly. The GamS version was constructed by around the horn PCR. Phage T7 Ocr was cloned into pCOLA (Novagen) by around the horn PCR on pUOS040 using primers listed in **Table S6**. All plasmids were transformed into *E. coli* Dh5α, extracted using the GeneJET Plasmid Miniprep kit (Thermo Fisher), verified by Sanger sequencing (Eurofins Genomics) and transformed into electrocompetent *E. coli* BL21-AI. Plasmids built for this study are listed in **Table S7**.

### Plaque assays

Phages were ten-fold serially diluted in LB and spotted on soft agar plates (0.7 % (w/v) agar) of *E. coli* BL21-AI containing a control plasmid or individual Kiwa systems. For spot assays in strains expressing phage proteins *in trans*, induction was performed with 0.2% or 0.02% L-arabinose. Plates were left to dry at room temperature and incubated overnight at 37 °C. Plaque forming units (PFUs) were determined by counting the plaques after overnight incubation. Whenever plaques were too small to be counted, a clear lysis area was set to be equal to 1 PFU or indicated by an open dot on the bar graph. Fold defence was determined by calculating the ratio of phage concentration of Kiwa expressing strain relative to control strains.

### Time post infection assays

Overnight bacterial cultures of *E. coli* BL21-AI with a control plasmid or individual Kiwa systems were diluted 1:100 in LB medium and grown at 37 °C with agitation until an OD_600_ of ≈0.3. Cultures were normalized and infected with phage T4 at an MOI of 3. A sample was taken at different time points post infection, serially diluted, and total phages were quantified by plaque assay on a phage-sensitive bacterial lawn (*E. coli* BL21-AI + pUOS016). For colony forming units, samples were washed twice with PBS 1x, serially diluted and spotted onto LBA plates. PFUs and CFUs were counted after overnight incubation at 37 °C.

### Conjugation assays

S17 cells containing plasmid pSEVA637 (donor) and BL21-AI cells containing the Kiwa systems or control plasmid (recipient) were grown overnight with antibiotics (gentamycin and chloramphenicol, respectively). Recipient cells were diluted 1:2 in LB and grown for at least 3h, while donor cells were diluted 1:100 in LB and grown until an OD_600_ of 0.3 at 37 °C, 180 rpm. 750 μl of donor cells were mixed with 250 μl of recipient cells, centrifuged at 9000 × *g* for 5 min, and resuspended in 25 μl of LB. The cell mixture was spotted onto LBA plates and incubated overnight at 30 °C. The cells were recovered from the plates and resuspended in 1 ml of PBS 1x. Two or ten-fold dilutions of this cell suspension were spotted onto LBA plates supplemented with chloramphenicol (for total recipient cell count) and LBA plates supplemented with chloramphenicol and gentamycin (for conjugant cell count). The plates were incubated at 37 °C for 18-24h and cells counted. Conjugation efficiency was calculated as the ratio (%) of conjugants per total recipient cells.

### DNA replication assay

Overnight bacterial cultures of *E. coli* BL21-AI with a control plasmid or individual Kiwa systems were diluted 1:100 in LB medium and grown at 37 °C with agitation until cultures reached an OD_600_ of ≈0.3. Cultures were normalized and infected with phage T3 or T4 at an MOI of 3. At 0 and 30 minutes post infection, a 5 ml sample was taken and centrifuged (15,000 × *g*, 5 min, 4 °C). The cell pellets were snap frozen using liquid nitrogen and stored at -80 °C. Total DNA was extracted using the GeneJET Genomic DNA Isolation kit (Thermo Fisher), using the Gram-negative protocol. Samples were sequenced at SeqCenter (Pittsburgh, PA, USA), where sample libraries were prepared using the Illumina DNA Prep kit and IDT 10bp UDI indices, and sequenced on an Illumina NextSeq 2000, producing 2×151bp reads. Demultiplexing, quality control and adapter trimming was performed with bcl-convert (v3.9.3). Reads were aligned to *E. coli* BL21-AI and phage T3 or T4 reference genomes (CP047231, NC_003298 and NC_000866) using MiniMap2^51^ in Geneious prime 2023.0.1. Replication efficiency was determined by calculating the increase of phage genomes to the number of cells at time 0. The percentage was calculated by comparison of Kiwa infected cells to control cells over time.

### Bacterial two-hybrid assay

Expression plasmids were cloned by fusing the T18 or T25 fragments of adenylate cyclase (CyaA) of *Bordetella pertussis* to either end of KwaB or the Lambda Gam protein, using primers listed in **Table S6**. *E. coli* BTH101 cells (F-, *cya-99*, *araD139*, *galE15*, *galK16*, *rpsL1* (*Str r*), *hsdR2*, *mcrA1*, *mcrB1*) were co-transformed with pairs of T18 and T25 plasmids, or control plasmids containing the leucine zipper motif of GCN4. Co-transformants were grown in LB supplemented with IPTG (0.5 mM), Ampicillin (100 µg/mL) and Kanamycin (50 µg/mL) at 30 °C with agitation. Overnight cultures were then spotted on LB agar plates supplemented with IPTG (0.5 mM), 5-bromo-4-chloro-3-indolyl-β-D-galactopyranoside (X-Gal, 40 µg/mL) and antibiotics. Plates were incubated at 30°C up to 48 hours before imaging.

### RecBCD mutant assays

Strains with individual deleted RecBCD subunits were transformed by electroporation with a control or Kiwa plasmids and recovered on LBA supplemented with kanamycin (50 µg/mL), chloramphenicol (25 µg/mL) and spectinomycin (50 µg/mL). Colonies were picked and grown overnight at 37 °C. Spot assays were performed with phages T4, T4Δgp2, Lambda_vir_, and Lambda_vir_ with a frameshift mutation in Gam.

### Isolation and sequencing of escape phages

Escape phages were isolated by picking single plaques at the lowest dilution of a spot assay on a lawn of *E. coli* cells expression Kiwa. The resistant phenotype of each escape phage was determined by comparing the EOP of the escape phage to the EOP of the wild type phage in the presence of Kiwa. Isolated phages were further propagated by infecting a liquid culture of Kiwa as described above. Phage lysates were pelleted following the NaCl/PEG8000 precipitation protocol as previously described^52^. The phage pellets were re-suspended in LB, and treated with DNase I and RNase A (1 µg/mL each) for 30 min at room temperature. EDTA (20 mM), proteinase K (50 µg/mL) and SDS (0.5%) were added to the suspension and incubated overnight at 56 °C. DNA was extracted using the Zymo Clean & Concentrator kit (Zymo Research). Samples were sequenced at SeqCenter (Pittsburgh, PA, USA), where sample libraries were prepared using the Illumina DNA Prep kit and IDT 10bp UDI indices, and sequenced on an Illumina NextSeq 2000, producing 2×151bp reads. Demultiplexing, quality control and adapter trimming was performed with bcl-convert (v3.9.3). Mutations were identified by mapping sequencing reads to the respective reference phage genomes (NC_003298 for T3 and **Table S3** for T7s) using Breseq^53^ v0.37.0 with default parameters. Only mutations that occurred in the isolated mutant but not in the wild type phage were considered. Silent mutations within protein-coding regions were also neglected.

### Amplification of mutant phages in cells expressing wild type tail fibres

Escape mutant T3 phages were propagated in cells expressing wild type T3p48 in *trans* for 4h at 37 °C, 180 rpm. The cultures were centrifuged at 9,000 × *g*, the supernatant recovered, filtered (0.2 μm PES), and stored at 4 °C until further use. The recovered phages were ten-fold diluted and spotted onto double layer agar with control or Kiwa-expressing cells. The plates were incubated overnight at 37 °C for determining phage titres.

### Protein expression and purification

Overnight cultures of *E. coli* BL21-AI with N-terminal 9xHis KwaB from ECOR8 or ECOR12 were diluted in LB supplemented with antibiotics, and grown at 37 °C with agitation until an OD_600_ of ≈0.6. Grown cultures were incubated on ice for 1 hour and protein expression was induced with L-arabinose (0.2 %) and IPTG (1 mM), followed by overnight incubation at 20 °C and 150 rpm. The overnight cultures were harvested by centrifugation (8,000 × g, 30 min, 4 °C). The supernatant was discarded and the cell pellets were re-suspended in lysis/wash buffer (50 mL per 1 L of initial culture, 100 mM Tris-HCl, 500 mM NaCl, 1 mM DTT, 5% glycerol, 40 mM imidazole, pH 8.0) supplemented with Pierce™ Protease Inhibitor Tablets EDTA-free (Thermo Fisher). Cell lysis was performed three times in a cooled French press (15Gpa). The cell lysate was centrifuged (17,000 × g, 30 min, 4 °C) and debris were removed by filtration (0.45 µm PES). The filtered supernatant was incubated with HisPur™ Ni-NTA Resin (Thermo Fisher) (500 µL/50 mL lysate) pre-washed with 20 mL of lysis/wash buffer for 30 min at 4 °C. The sample was loaded onto a 5 mL Pierce™ Disposable Column (Thermo Fisher) for gravity-flow affinity chromatography. The column was washed with 15 mL of ice-cold lysis/wash buffer, and the protein was eluted with ice-cold elution buffer (100 mM Tris-HCl, 500 mM NaCl, 1 mM DTT, 5% glycerol, 250 mM imidazole, pH 8.0). Pooled fractions were concentrated (Amicon Centrifugal concentrator with Ultracel membrane, 10 kDa MWCO), and the buffer exchanged to running buffer (100 mM Tris-HCl, 150 mM NaCl, 1 mM DTT, 5% glycerol, pH 8.0). The resulting protein sample was subjected to size exclusion chromatography using a Superdex 200 increase 10/300 GL (Cytiva) column equilibrated with running buffer with 0.5 mL/min flow rate using running buffer as a mobile phase. Fractions of interest were analysed by SDS-PAGE, snap frozen in liquid nitrogen and stored at -80 °C until further use.

### KwaA expression and purification

The genes encoding KwaA (UniProt: P0DW45) and KwaB (UniProt: P0DW46) from *Escherichia coli* O55 were codon-optimized and synthesized by IDT. They were then individually cloned into the pET21a vector (Novagen) with ampicillin resistance and the pYB100 vector (NovoPro) with kanamycin resistance. KwaA was fused to a C-terminal Flag tag, while KwaB was fused to a C-terminal His6 tag.

The O55 KwaA membrane protein was overexpressed in C43 cells, a derivative of *Escherichia coli* BL21(DE3), and induction was carried out using 0.5 mM IPTG (GoldBio) at 16 °C for 20 hours. The cells were harvested by centrifugation and resuspended in a lysis buffer (25 mM Tris, pH 8.0, 500 mM NaCl, 2 mM β-mercaptoethanol, and cOmplete (EDTA-free protease inhibitor). Cell lysis was achieved through sonication, after which 1% (w/v) Lauryl Maltose Neopentyl Glycol (LMNG, Anatrace) and 0.1% (w/v) cholesteryl hemisuccinate (CHS, Anatrace) were added to solubilize the proteins at 4 °C for 3 hours. Insoluble material was cleared by centrifuging at 22,000 rpm for 1 hour using a JA-20 fixed-angle rotor (Avanti J-E series centrifuge, Beckman Coulter). The resulting supernatant was incubated with anti-FLAG M2 affinity gel (SIGMA, A2220) while rotating at 4 °C for another 3 hours. The sample was subsequently loaded onto a gravity column and thoroughly washed with Wash Buffer I (25 mM Tris, pH 8.0, 500 mM NaCl, 2 mM β-mercaptoethanol, 0.1% (w/v) LMNG, and 0.01% (w/v) CHS), followed by Wash Buffer II (25 mM Tris, pH 8.0, 500 mM NaCl, 2 mM β-mercaptoethanol, 0.001% (w/v) LMNG, 0.0001% (w/v) CHS, and 0.00033% Glyco-diosgenin (GDN, Anatrace)). The protein was then eluted with Wash Buffer II containing 0.2 mg/ml 3× DYKDDDDK Peptide (Pierce) while rotating for 30 minutes at 4 °C. The elution was then concentrated, and further purified using size exclusion chromatography (SEC) on a Superose 200 Increase 10/300 GL column (GE Healthcare) equilibrated with SEC buffer (25 mM Tris, pH 8.0, 500 mM NaCl, 2 mM β-mercaptoethanol, 0.001% (w/v) LMNG, 0.0001% (w/v) CHS, and 0.00033% GDN). Fractions eluted at around 15 ml, indicating tetrameric assembly, were collected, concentrated, and prepared for SDS-PAGE and cryo-EM analysis.

### KwaB expression and purification

KwaB protein purification for crystallization began with its expression in BL21 (DE3) cells, using a C-terminal His tag. Protein expression was induced with 0.5 mM IPTG, and the culture was grown at 16 °C with shaking at 220 rpm for about 20 hours. The cells were then harvested by centrifugation at 4,000 rpm for 15 minutes, and the pellet was resuspended in a lysis buffer containing 25 mM Tris (pH 8.0), 500 mM NaCl, 5% glycerol, 2 mM β-mercaptoethanol, and PMSF. After cell disruption using sonication, the lysate was centrifuged at 22,000 rpm for 1 hour to separate the soluble fraction. The clarified lysate was then applied to a Ni-NTA gravity column, where the protein underwent flow-through, washing, and elution steps. The eluted protein solution contained 25 mM Tris (pH 8.0), 150 mM NaCl, 5% glycerol, 2 mM β-mercaptoethanol, and 500 mM imidazole. To further purify the sample and remove any nucleic acid contaminants, it was passed through a heparin column. Finally, the protein underwent gel filtration using a Superose 200 Increase column to exchange it into the crystallization buffer, which contained 25 mM HEPES (pH 7.5), 150 mM NaCl, 5% glycerol, and 2 mM DTT.

### KwaAB expression and purification

For cryo-EM analysis of the KwaA and KwaB complex, KwaA_flag and KwaB-His6 were co-expressed and purified using anti-FLAG resin, following the same protocol as that used for isolating KwaA alone. The fraction peaks from the Superose 6 Increase column revealed the high molecular weight KwaAB complex at 11.8 ml and the low molecular weight KwaAB complex at 14.5 ml, as indicated by the SDS-PAGE results. Additionally, the MscK protein co-eluted with the high molecular weight complex, which was also confirmed by SDS-PAGE analysis.

### Cryo-EM sample preparation and data collection

For membrane bound KwaA, the purified protein was concentrated to 4 mg/ml using a Superose 6 Increase column. A 4 µl aliquot of the sample was applied to glow-discharged holey gold grids (UltrAuFoil 300 mesh R1.2/1.3). The grids were blotted for 3 seconds with a force setting of 0 at 6 °C and 100% humidity, then rapidly frozen in liquid ethane using a Vitrobot Mark IV (FEI). Data collection was performed at the Memorial Sloan Kettering Cancer Center (MSKCC) using a Titan Krios G2 transmission electron microscope (FEI) operating at 300 kV and equipped with a K3 direct detector, controlled by SerialEM software. Movies were recorded in super-resolution mode with a total electron dose of 53 e^−^/Å², a defocus range of -0.8 to -2.2 µm, and a pixel size of 1.064 Å. The cryo-EM sample preparation for the KwaAB complex was performed similarly to that of KwaA alone. The peak corresponding to the high molecular weight complex from the Superose 6 Increase column was collected and concentrated to 4.5 mg/ml for cryo-EM analysis. Data collection was carried out using a Titan G2 transmission electron microscope (FEI) operating at 300 kV, equipped with a K3 electron detector, and controlled by Leginon software at the New York Structural Biology Center (NYSBC). The defocus range was set between - 0.8 to -2.2 µm, with a pixel size of 0.826 Å and a total electron dose of 58.73 e^−^/Å² in super-resolution mode.

### Cryo-EM data processing

For KwaA structure determination, a total of 6,054 movies was processed using cryoSPARC^54^, Patch motion correction and patch contrast transfer function (CTF) estimation were applied to correct for drift and estimate CTF parameters, respectively. Micrographs with ice contamination, high astigmatism, or poor CTF fit resolution were excluded by setting appropriate threshold ranges using the ’Manually Curate Exposures’ job. High-quality micrographs were then processed with the Blob picker and extract jobs, resulting in the selection of 5,479,293 particles. Multiple rounds of two-dimensional (2D) classification were conducted to eliminate junk particles. After two rounds of ab-initio reconstruction and heterogeneous refinement into three classes, two distinct volumes with C2 and C4 symmetries were identified. These were then refined further through homogeneous and non-uniform refinement. For the C2 symmetry reconstruction, 312,208 particles were used, achieving a resolution of 3.74 Å, which enabled precise model building. In the case of C4 symmetry reconstruction, 158,542 particles were utilized, reaching a resolution of 4.29 Å, sufficient for fitting the main chain.

To determine the structure of the KwaAB complex, 5,475 micrographs were collected using the K3 camera, and the movies were processed using the same procedure. Following particle selection through Blob picking and template-based methods, 1,731,427 particles were chosen for 2D classification. From these 2D images, the KwaA-KwaB complex and Msck protein were separated, resulting in 182,763 particles for the KwaAB complex and 90,497 particles for the MscK protein. The particles from the KwaA-KwaB complex were subjected to ab-initio reconstruction to generate initial models, followed by heterogeneous refinement to enhance the density map’s quality. The best class from the heterogeneous refinement was selected and further polished using homogeneous and non-uniform refinement, achieving a 3.75 Å resolution structure with C4 symmetry. This level of detail allowed for accurate model building of the KwaA tetramer bound to KwaB dimers. For the MscK protein group, the same protocol as the KwaAB complex was followed, achieving a resolution of 4.14 Å, which enabled fitting the MscK heptamer in its closed state. All the resolution estimations were based on a Fourier shell correction of 0.143 cutoff.

### Crystallisation and data collection

KwaB dimer crystals were obtained at 20 °C using the hanging-drop vapor diffusion technique, where 1 µL of the protein solution was combined with 1 µL of reservoir solution containing 0.1 M Tris (pH 8.5) and 5% (w/v) PEG8000. The crystals were cryoprotected by adding 25% glycerol to the reservoir solution before being flash cooled in liquid nitrogen. Diffraction data were collected at the FMX beamline of NSLS-II at Brookhaven National Laboratory and processed using the HKL suite (HKL Research).

### Model building, structure refinement and visualization

The KwaA tetramer and KwaB dimer models were predicted using Alphafold2^55^, while the MscK model was based on the published heptamer structure (PDB: 7UW5)^56^. The KwaAB model was constructed by fitting the KwaA tetramer and KwaB dimer separately, followed by structure building in Coot^57^. For KwaB crystals, the structure was solved by molecular replacement using PHASER^58^, all of the atomic coordinates were refined against the map in PHENIX^59^.Figures were prepared using UCSF Chimera^60^ and USCF ChimeraX^61^ with the final image layout created in Adobe Photoshop and Illustrator.

### KwaB mutants’ expression and purification

KwaB mutants targeting the DNA-binding site were designed based on the AF3 model. Two mutants, named KwaB 3-mut (R142A/R167A/K187A) and KwaB 6-mut (R142A/R167A/K187A/R233A/N134A/D169A), were constructed as follows: the genes were codon-optimized and synthesized using IDT gBlock fragments, then cloned into the pYB100 vector using the Gibson cloning kit. Expression and purification of the mutants followed the protocol described above for crystallisation of the KwaB wild type protein. The purified proteins were concentrated to 2 mg/ml for subsequent functional assays.

### Electrophoretic mobility shift assay

For the O55 KwaB binding assays with ssDNA or dsDNA, we utilized a 72 nt sequence (5’TGGTTTTTATATGTTTTGTTATGTATTGTTTATTTTCCCTTTAATTTTAGGATATGAAAACAAGAATTTATC) as well as its complementary DNA substrates. The DNA substrates were assembled through self-annealing in an annealing buffer containing 25 mM HEPES (pH 7.5), 150 mM NaCl, and 2 mM MgCl₂.For the assembly of DNA with KwaB and mutant KwaB proteins, 10 nM DNA substrates were incorporated into a total volume of 20 µl. The ratios of KwaB protein to dsDNA were set at 0, 1, 8, 16, 32, 64, and 128:1, while for ssDNA, the ratios were 0, 1, 4, 8, 16, 32, and 64:1. The protein and DNA substrates were diluted in an annealing buffer that contained an additional 5% glycerol for gel loading. After incubating the mixtures on ice for 1 hour, 10 µl of each was loaded onto a 6% DNA Retardation Gel (Invitrogen) and electrophoresed at 100 V for 35 minutes at 4 °C in 0.5 x TBE buffer. Nucleic acids were visualized using SYBR Gold nucleic acid gel stain (Invitrogen), incubated in the dark for 30 minutes, and imaged using a ChemiDoc imaging system.

Additionally, pUC19 plasmid DNA, *E. coli* BL21-AI gDNA, and phage Lambda gDNA (Thermo Fisher) were used as DNA probes for EMSA assays. 50 ng of dsDNA probe or 150 ng ssDNA (FN0850) were mixed with purified KwaB in assembly buffer 1 (100 mM Tris-HCl, 150 mM NaCl, 1 mM DTT, 5% glycerol, 10 mM MgCl_2_, pH 8.0) and incubated at 37 °C for 30 minutes. For titrations, dilutions ranging from 1400 to 11 ng of KwaB protein were mixed with the DNA probe prior to complex assembly.

Samples were mixed with blue DNA loading dye (without EDTA) (Thermo Fisher) and loaded on a 1% agarose gel for electrophoresis. Gels were run at 100 V for 50 minutes in 1X TAE buffer. Nucleic acids were visualised with SYBRSafe or SYBRGold staining and captured with an Invitrogen iBright 1500 apparatus (Thermo Fisher).

To assess the impact of mutations in residues involved in the KwaB-DNA binding surface, purified KwaB wild type (wt) protein and mutants (3-mut and 6-mut) were concentrated to 2 mg/ml. A 128:1 ratio of KwaB proteins to DNA substrate was used, with 50 ng of 72-nt single-stranded DNA (ssDNA) or 72-bp double-stranded DNA (dsDNA) added to a total reaction volume of 10 μl. The mixtures were incubated at room temperature for 30 minutes. Subsequently, 3 μl of each sample was loaded onto a 6% DNA retardation gel (12-well), along with 0.2 μl of Quick-Load Purple Low Molecular Weight DNA Ladder (NEB, N0557S). The gel was run at 100 V for 80 minutes at 4°C in 0.5× TBE running buffer. SYBR Gold was used to stain the gel in the dark with gentle shaking for 30 minutes, followed by three washes with ddH2O. The gel was imaged using a ChemiDoc imaging system.

### *In vitro* nuclease activity

Purified PCR amplicons, pUC19 plasmid DNA, *E. coli* BL21-AI gDNA, phage Lambda gDNA (Thermo Fisher) and oligos were used as substrates for cleavage assays. Target cleavage assays were performed in a 10 µl reaction mixture containing 50 ng dsDNA or 600 ng ssDNA (FN0850) substrate and 500 ng protein in cleavage buffer (100 mM Tris-HCl, 150 mM NaCl, 1 mM DTT, 5% glycerol, pH 8.0) supplemented with ddH_2_0, MgCl_2_ (10 mM) or MnCl_2_ (10 mM), and ATP (1mM). Assays were allowed to proceed at 37 °C for 2 hours and stopped by addition of proteinase K (Thermo Fisher) and incubated at 55 °C for 15 minutes. Samples were mixed with purple DNA loading dye (with EDTA) (NEB) and loaded on a 1% or 1.5% (oligos only) agarose gel for electrophoresis. Gels were run at 100 V for 50 minutes in 1X TAE buffer. Nucleic acids were visualised with SYBRSafe or SYBRGold (oligos only) staining and captured with an Invitrogen iBright 1500 apparatus (Thermo Fisher).

### Fluorescence anisotropy binding assays

To determine DNA binding affinities of KwaB, serial two-fold dilutions of ECOR8 KwaB were incubated with 0.8 nM FAM-labelled 20-nt ssDNA (top oligo, **Table S6**) or 20-bp dsDNA (top and bottom oligo, **Table S6**) for 3 hours at 37 °C in binding buffer (150 mM NaCl, 25 mM HEPES pH 7.5, 0.05 Tween-20). Data were recorded at 37 °C in a CLARIOstar Plus multi-detection plate reader (BMG Labtech) equipped with a fluorescence polarization optical module (λ_ex_ = 485 nm; λ_em_ = 520 nm). Data were fit in GraphPad Prism 10 using a one-step binding model, and extrapolated start and end points were used to normalize the calculation of fraction bound. For Gam competition binding experiments, 1 µM KwaB was preincubated with 50 nM ss/dsDNA as described above, and then further supplemented with a final concentration of 10 µM GamS (New England Biolabs), and then measurements were performed as described above.

### Pull-down assay for GamS and KwaB

The full-length GamS was fused to a C-terminal Flag tag and cloned into the pET21a vector. It was then co-transfected into *E. coli* with the KwaB-His plasmid at a 4:1 molar ratio, followed by selection based on ampicillin and kanamycin resistance.The selected colonies were then transferred to 1 L of LB liquid medium, induced, and harvested using the same method. The bacteria were lysed with Lysis buffer (25 mM Tris, PH 8.0, 300 mM NaCl, 5% glycerol, 2 mM DTT), followed by low-temperature sonication and centrifugation to obtain the supernatant. This supernatant was incubated with Flag beads for 3 hours, after which the beads were washed, and the target protein was eluted using 3x Flag peptide. The eluate was concentrated and loaded onto Superose 6 Increase column, and peaks 2 and 3 were collected for gel analysis. SDS-PAGE results demonstrated the co-elution of KwaB and Gam.

### Confocal microscopy

Overnight cultures of control (YFP) or Kiwa-expressing cells were sub-cultured in LB with antibiotics for 2 hours. Agarose pads were prepared with 1% agarose in LB. For membrane staining, cells were washed with 1× PBS and resuspended in 0.1 mM FM4-64 (in PBS). Two μl of bacterial culture were seeded onto the agarose pads and covered with microscope cover slips prior to imaging. Confocal images and movies were acquired using a dual point-scanning Nikon A1R-si microscope equipped with a PInano Piezo stage (MCL), using a 60x PlanApo VC oil objective NA 1.40. Movies and images were acquired in galvanometer scanning mode using 488nm Laser (Coherent, 50mW), and a 561 nm Laser (Coherent, 50mW). Image processing and quantification was performed using FiJi software. The phage filtrates were treated with DNase I (1 µg/ml each) for an hour at room temperature. SYTOX orange was added to a final concentration of 2.5 µM, and incubated overnight at 4 °C. Excess dye was removed by centrifuging the stained lysate on a 100 kDa Amicon Ultra (UFC5100) filter and rinsing the dye away with fresh LB four times. The concentrated stained phages were resuspended in 1 ml of fresh LB and stored at 4 °C.

### RNAseq

Overnight cultures were diluted with fresh LB supplemented with antibiotics at 1:100 and incubated at 37 °C, 180 rpm for 2 hours. Phages were added to the cells at MOI 10. The cells were harvested by centrifugation after 15 minutes of incubation, and the RNA immediately extracted with RNeasy Kit (Qiagen). Library construction, rRNA depletion, and paired-end Illumina sequencing (Novaseq 6,000, 2 × 150 bp configuration) were performed by Novogene.

### Bioinformatic analysis

Information on defence systems in 223,143 prokaryotic genomes was retrieved from the PADLOC database^13,14^. Only complete Kiwa systems that contained both KwaA and KwaB gene within the operon, were considered for further analysis (**Table S8**). For phylogenetic analyses, split KwaB proteins were concatenated into a single sequence, and KwaA and KwaB proteins were dereplicated using MMseqs2 at 90% sequence identity threshold^62^. Contigs encoding representative KwaA and KwaB proteins shorter than 150 and 200 amino acids, respectively, were re-examined. Partial sequences located at the ends of contigs were discarded, and any sequences with updated annotations in the latest RefSeq release (v227) were replaced with the new annotations. The retained sequences (**Tables S9, S10**) were aligned using Muscle5^63^, and sites with more than 99% of gaps were trimmed using ClipKIT^64^. Phylogenetic trees were reconstructed using IQ-Tree2^65^, and the optimal model was picked using ModelFinder from the set of WAG, LG, Q.Pfam, and NQ.Pfam^66^. Support values were estimated using 1,000 ultrafast bootstrap replicates with a hill-climbing nearest neighbour interchange optimization (UFBoot2)^67^, and trees were visualized using iTOL v6^68^. Contigs encoding KwaA and KwaB from the dereplicated set were searched for plasmids and prophages using geNomad v1.7^69^, and genomic islands were identified using TreasureIsland with probability cutoff at 0.95^70^. Additionally, 10 genes upstream and downstream of KwaA and KwaB were annotated using COG profiles^17^. Kiwa genes that are colocalized with at least 2 hallmark genes from MGEs (X category) were labelled as associated with ‘other mobilome’ (**Tables S9, S10**).

Protein domains of KwaA and KwaB were identified using HHpred^71^. Transmembrane domains were identified using DeepTMHMM^72^. KwaA proteins with more or less than 4 transmembrane domains were manually checked for misannotation of the start codon. Sequence alignments of KwaA C-terminus and KwaB were constructed using Clustal Omega^73^ and visualised using Geneious Prime v2023.0.1. For KwaB mutants, a total of nine homologs present in the KwaB ECOR8 and ECOR12 cluster were selected and aligned to determine amino acid conservation. KwaA schematic was visualised and adapted from Protter^74^. All structural models were built using Alphafold^75^ (AF2) software run on the external COSMIC server^76^ using either the multimer or monomer model with complete database settings. Structural similarity was estimated using DALI^77^ for monomers, and MM-align^78^ for multimers. Structures were visualised using ChimeraX v1.6.1^79^. KwaB functionality was predicted using the Biomedical AI Platform webtool SPROF-GO^29^.

Genomes of *E. coli* BL-21-AI and T2 phage used in this study were annotated using PGAP pipeline^80^. The quality of RNA-seq data was assessed using FastQC, and low-quality tiles were removed using BBTools^81^. Reads were mapped to the genome using HISAT2^82^, and the ones corresponding to tRNA and rRNA were discarded. TPM values were calculated using TPMCalculator^83^, and for mathematical purposes, phage genes and bacterial genes were treated as separate datasets.

## QUANTIFICATION AND STATISTICAL ANALYSIS

Unless stated otherwise, experiments were performed in biological triplicates and are represented as the mean and standard deviation. Statistical significance was calculated by ratio paired t-test or by two-way ANOVA with sidak’s multiple comparison test, with a significance level of 0.05.

## Supplemental information titles and legends

**Figure S1** Similarity of Kiwa homologues and their anti-phage activity.

**Figure S2** Structure analysis of KwaA.

**Figure S3** Interactions between KwaB monomers in the symmetrical dimer.

**Figure S4** Structure analysis of KwaAB.

**Figure S5** Validation of phage staining with SYTOX Orange.

**Figure S6** KwaB does not degrade DNA but binding to phage DNA prevents replication.

**Figure S7** Inhibition of KwaB by Lambda Gam protein.

**Table S1** Cryo-EM data collection and refinement statistics.

**Table S2** Crystal data collection and refinement statistics (molecular replacement).

**Table S3** Mutation analysis of phage escape mutants. Mutations marked by Breseq under "marginal evidence" are reported if the mutation was supported by more than 20% of the aligned reads.

**Table S4** RNAseq data of Kiwa and control (YFP) cells infected with phage T2 for 15 min.

**Table S5** Strains and phages used in this study.

**Table S6** Primers and gblocks used in this study.

**Table S7** Plasmids used in this study.

**Table S8** Annotation of 9,797 Kiwa operons in 9,582 prokaryotic genomes based on PADLOC database.

**Table S9** Annotation of mobile genetic elements in the dereplicated dataset of KwaA genes.

**Table S10** Annotation of mobile genetic elements in the dereplicated dataset of KwaB genes.

