## Supplementary material for "Kiwa is a bacterial membrane-embedded defence supercomplex activated by phage-induced membrane changes": Figure S

### Contents

**Figure S1** Similarity of Kiwa homologues and their anti-phage activity.

**Figure S2** Structure analysis of KwaA.

**Figure S3** Interactions between KwaB monomers in the symmetrical dimer.

**Figure S4** Structure analysis of KwaAB.

**Figure S5** Validation of phage staining with SYTOX Orange.

**Figure S6** KwaB does not degrade DNA but binding to phage DNA prevents replication.

**Figure S7** Inhibition of KwaB by Lambda Gam protein.

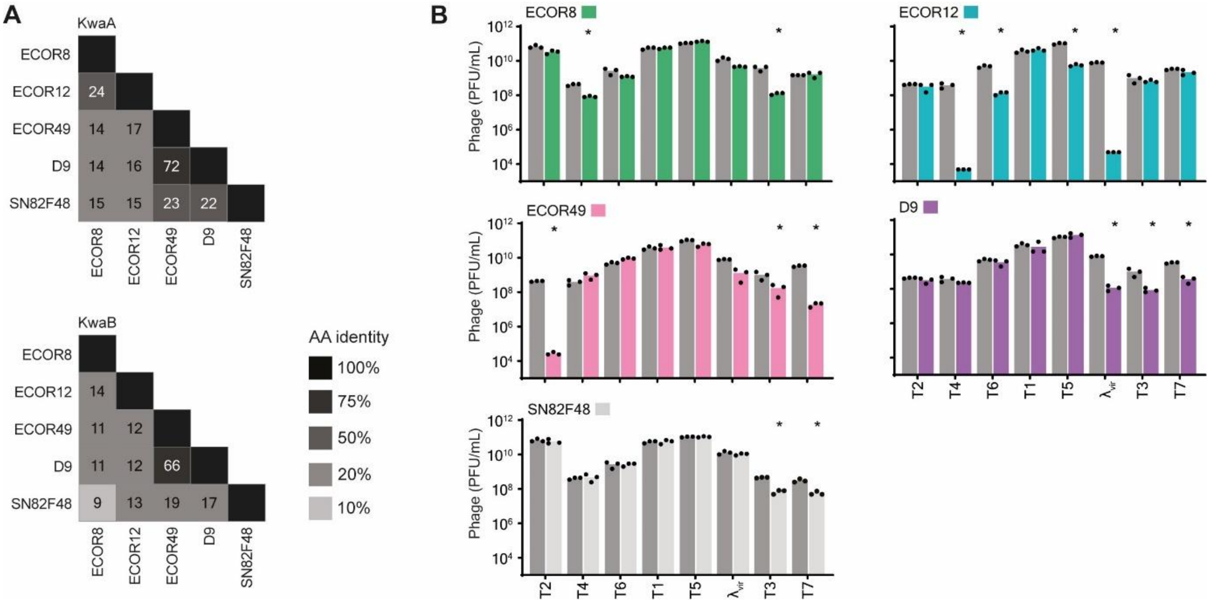


**Figure S1** Similarity of Kiwa homologues and their anti-phage activity.

**(A)** Relative amino acid identity of KwaA and KwaB from experimentally validated systems in this study.

**(B)** Data of plaque assays for all tested Kiwa systems shown in **Figure 1D**. Data represents phage PFU/mL on Kiwa systems (coloured bars) or control strains (grey). Bar graphs represent the average of three replicates, with individual data points overlaid. Open points indicate instances where it was not possible to determine individual phage plaques, hence a value of 1 was assumed.

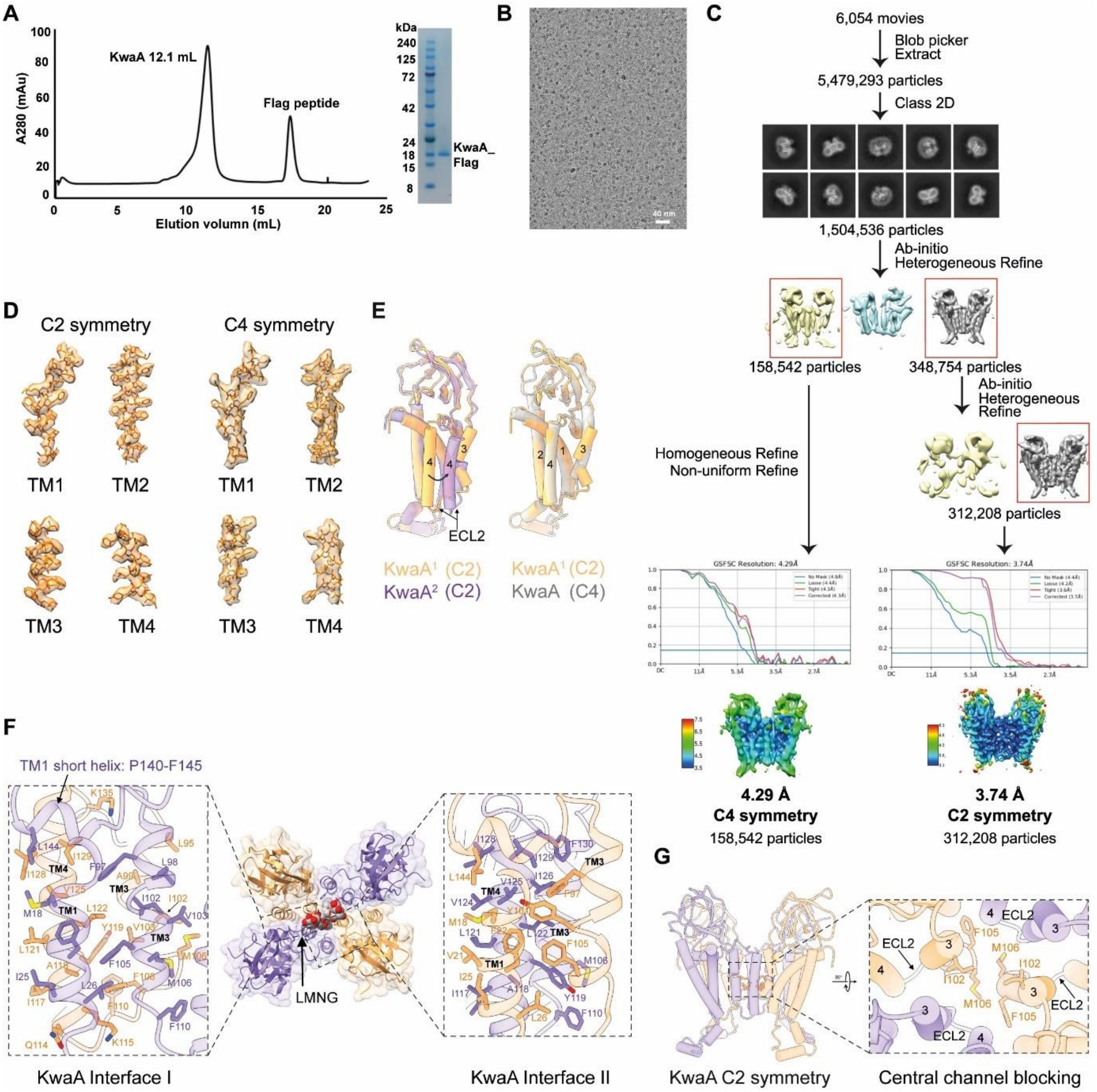


**Figure S2** Structure analysis of KwaA.

**(A)** Size exclusion chromatography (SEC) and SDS-PAGE analyses of the KwaA tetramer.

**(B)** A representative raw micrograph of the KwaA tetramer obtained by cryo-EM.

**(C)** Image processing flow of cryo-EM KwaA tetramer.

**(D)** Local density mapping of TM1-4 for both C2 and C4 symmetric KwaA structures.

**(E**) Structural alignment of two KwaA protomers. The comparison between KwaA protomers in the C2 symmetric tetrameric structure is shown in left panel, while the comparison between KwaA protomers in C2 and C4 tetrameric structures is shown in the right panel.

**(F)** KwaA exhibits two distinct interfaces within the C2 symmetry. The LMNG interface is indicated by an arrow in the central panel. On the left panel, detailed interactions at Interface I are represented. Key residues involved in these hydrophobic interactions from one protomer include M18, F22, I25, and L26 in TM1, and F97, L98, I102, V103, F105, M106, and F110 in TM3. In the adjacent protomer, the critical residues are L95, A99, I102, V103, M106, and F108 in TM3, and I117, A118, L121, L122, V125, I128, and I129 in TM4. Notably, F22 in TM1, along with F105, F110 in TM3 and F108, F110, Y119 in the neighbouring protomer form strong local π-π interactions. TM1 is skewed in the middle, where a short helix (residues P140-F145), positioned parallel to the cell membrane, establish hydrophobic interaction with TM4 via residue L144. Polar interactions are less prominent at interface I, with only residues Q114, K115 and K135 perhaps establishing the interaction with the main chain. On the right panel, detailed interactions at Interface II are shown, with hydrophobic and π-π interactions as observed in interface I.

**(G)** C2 symmetric KwaA tetramer with the expanded box showing alignment of residues lining the central channel of the tetramer. Residues I102, F105 and M106 from both protomers form a hydrophobic core that blocks the central channel.

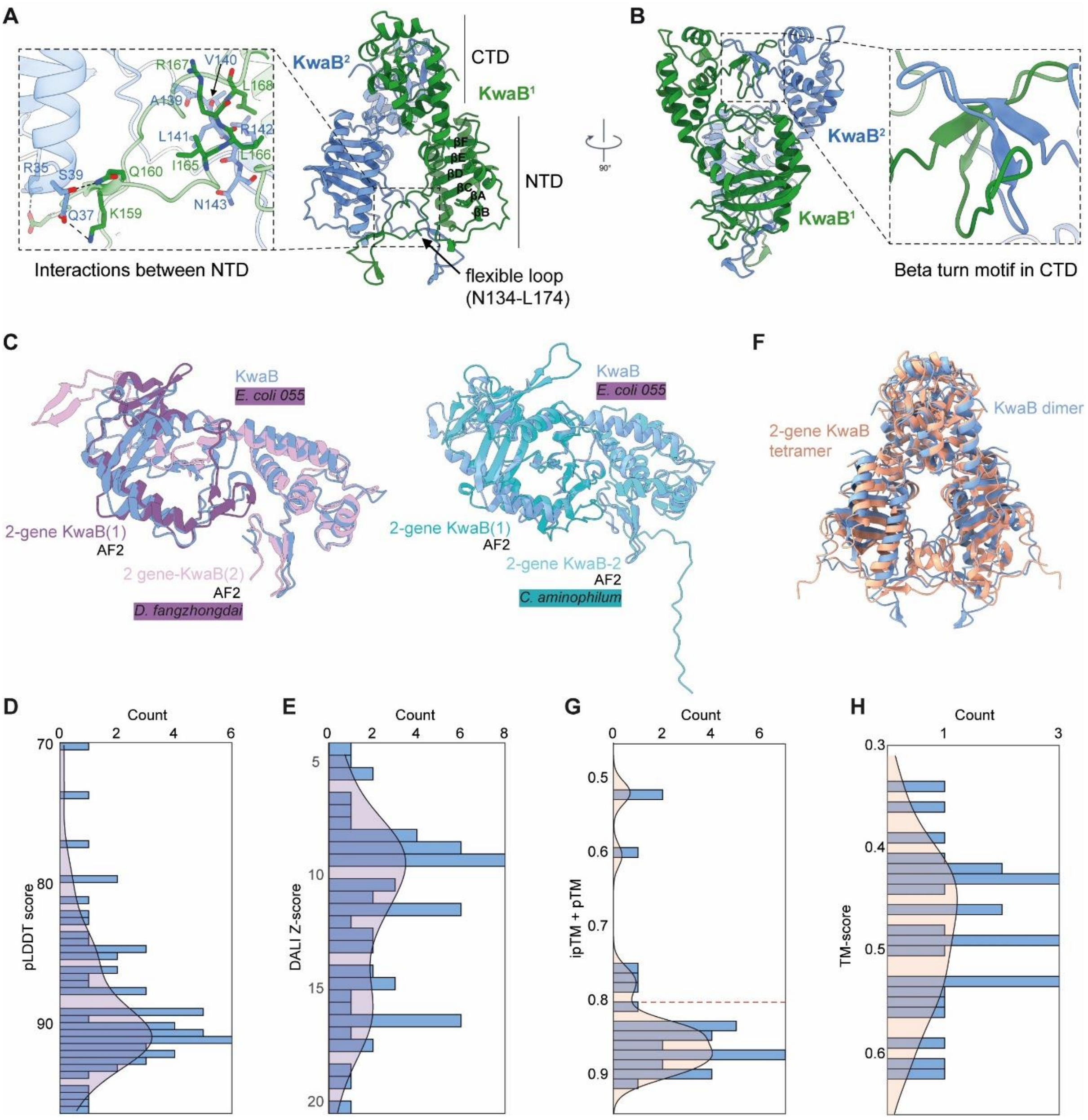


**Figure S3** Interactions between KwaB monomers in the symmetrical dimer.

**(A)** Interactions between N-terminal domains of KwaB monomers in the symmetrical dimer. The interactions in the expanded box of the N-terminal segment show that a flexible loop (134-174) between β-sheets D and E (DE loop) plays a crucial role in stabilizing the dimer interface. This loop is sandwiched between the identical regions and the N-terminal domain of the opposing KwaB protomer, forming strong interactions with both sides. Specifically, residues A139-N143 from one monomer align anti-parallel with residues I165-D169 from the other monomer, establishing multiple main-chain interactions. On the opposite side, D155 and K159 probably form a pair of salt bridge with R35 and E37, respectively. Additionally, at the periphery, Q160 potentially interacts with S39 through a hydrogen bond, further stabilizing the interface.

**(B)** Interactions between C-terminal domains of KwaB monomers in the symmetrical dimer. The interactions in the expanded box of the C-terminal segment show a beta-turn motif (similar to DE loop in N-terminus) that is also sandwiched between the same regions and the extreme C-terminal tail of the other KwaB protomer. The parallel binding interfaces in both N- and C-terminal regions indicate a tightly packed dimer structure, enhancing the stability of the KwaB homodimer.

**(C)** Superimposition of KwaB from *E. coli* O55 with the AlphaFold 2 predicted structures of the 2-gene KwaB proteins from *Dickeya fangzhongdai* (WP_225622576.1 and WP_225622577.1) (left) and *Clostridium aminophilum* (WP_074650070.1 and WP_074650071.1) (right). The KwaB containing strains are coloured by phylogenetic cluster as in Figure 1A.

**(D,E)** pLDDT score (D) and DALI Z-score (E) of the structures shown in (C).

**(F)** Superimposition of the KwaB dimer from *E. coli* O55 with the AlphaFold 2 predicted structure of the 2-gene KwaB tetramer from *Acinetobacter portensis* (WP_163122822.1 and WP_163122825.1).

**(G,H)** ipTM+pTM (G) and TM scores (H) of the complex shown in (F).

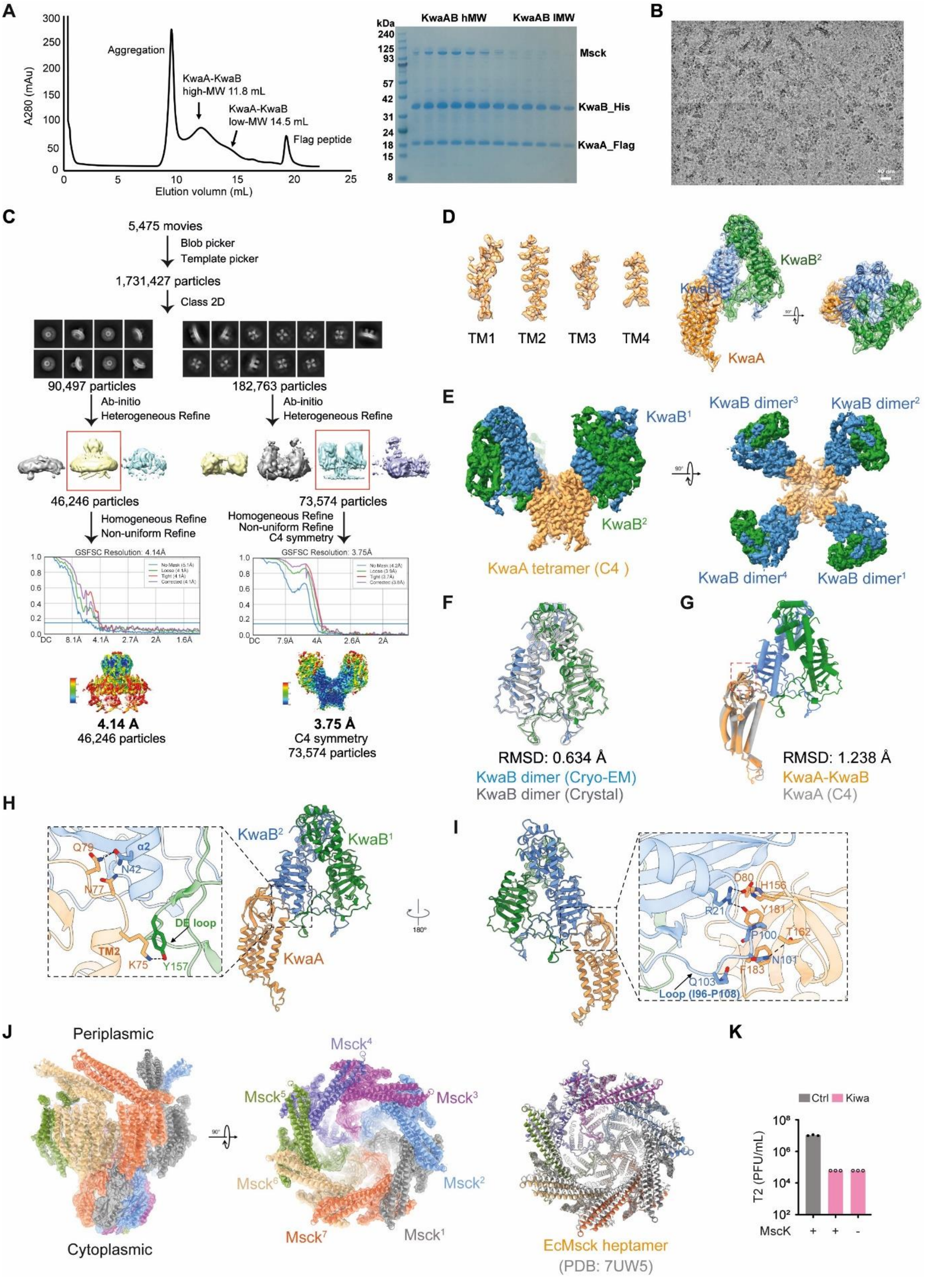


**Figure S4** Structure analysis of KwaAB.

**(A)** SEC and SDS-PAGE results demonstrated successful co-expression of the KwaAB complex, revealing both high and low molecular weight forms. SDS-PAGE indicated Msck was co-eluted with the high molecular weight fraction.

**(B)** A representative raw micrograph of higher molecular weight KwaAB complex.

**(C)** Image processing workflow of the cryo-EM structure of the high molecular weight KwaAB complex. A 4.1 Å map for the MscK protein was also generated from this dataset.

**(D)** Local density mapping of TM1-4 for KwaA, including the map fitting of the KwaA monomer and KwaB dimer.

**(E)** Two views of the cryo-EM density map of the high molecular weight KwaAB complex.

**(F)** KwaB dimer obtained from cryo-EM structure of the high molecular weight KwaAB and the crystal structure of KwaB exhibit an RMSD of 0.634 Å..

**(G)** Structural alignment of the KwaAB dimer in the high molecular weight KwaAB complex with the KwaA protomer exhibiting C4 symmetry, resulting in an RMSD of 1.238 Å.

**(H, I)** In-depth interactions between the KwaA monomer and KwaB dimers in the high molecular weight KwaAB complex. The smaller and larger interfaces are shown in the blowup boxes in panels H and I, respectively.

**(J)** Map fitting of the *E. coli* MscK heptamer (left and middle). The MscK protein displayed a closed state in the sample of high molecular weight KwaAB complex. The *E. coli* MscK structure closely matches the MscK closed state structure (7UW5) in the periplasmic gating ring.

**(K)** Anti-phage activity of Kiwa in *E. coli* BW25113 wild-type (control) or with deleted MscK. Data represents phage PFU/mL on tested strains. Asterisks indicate a statistically significant decrease in PFU/ml compared to control (Two-way ANOVA, $p < 0.05$). The bars represent the average of three biological replicates with individual data points overlaid.

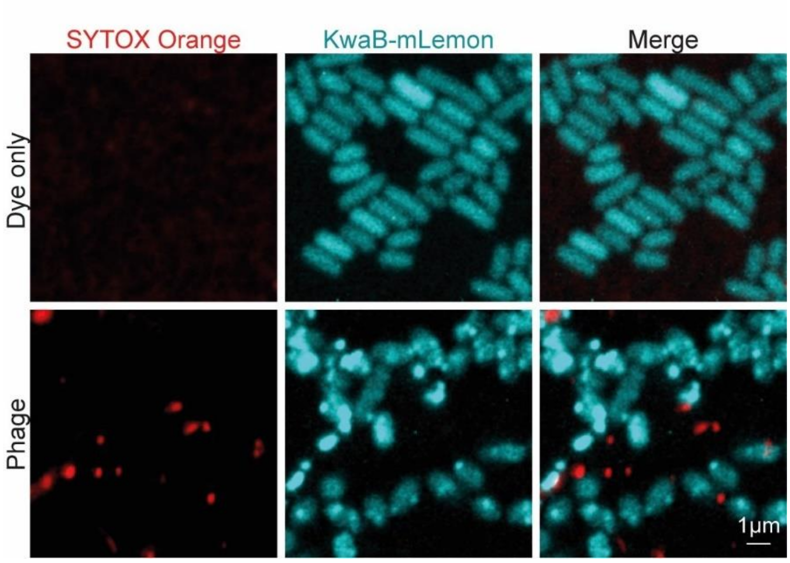


**Figure S5** Validation of phage staining with SYTOX Orange.

Confocal microscopy images of Kiwa-expressing cells exposed to SYTOX Orange or phage stained with SYTOX orange. Red dots are only observed when stained phage is present.

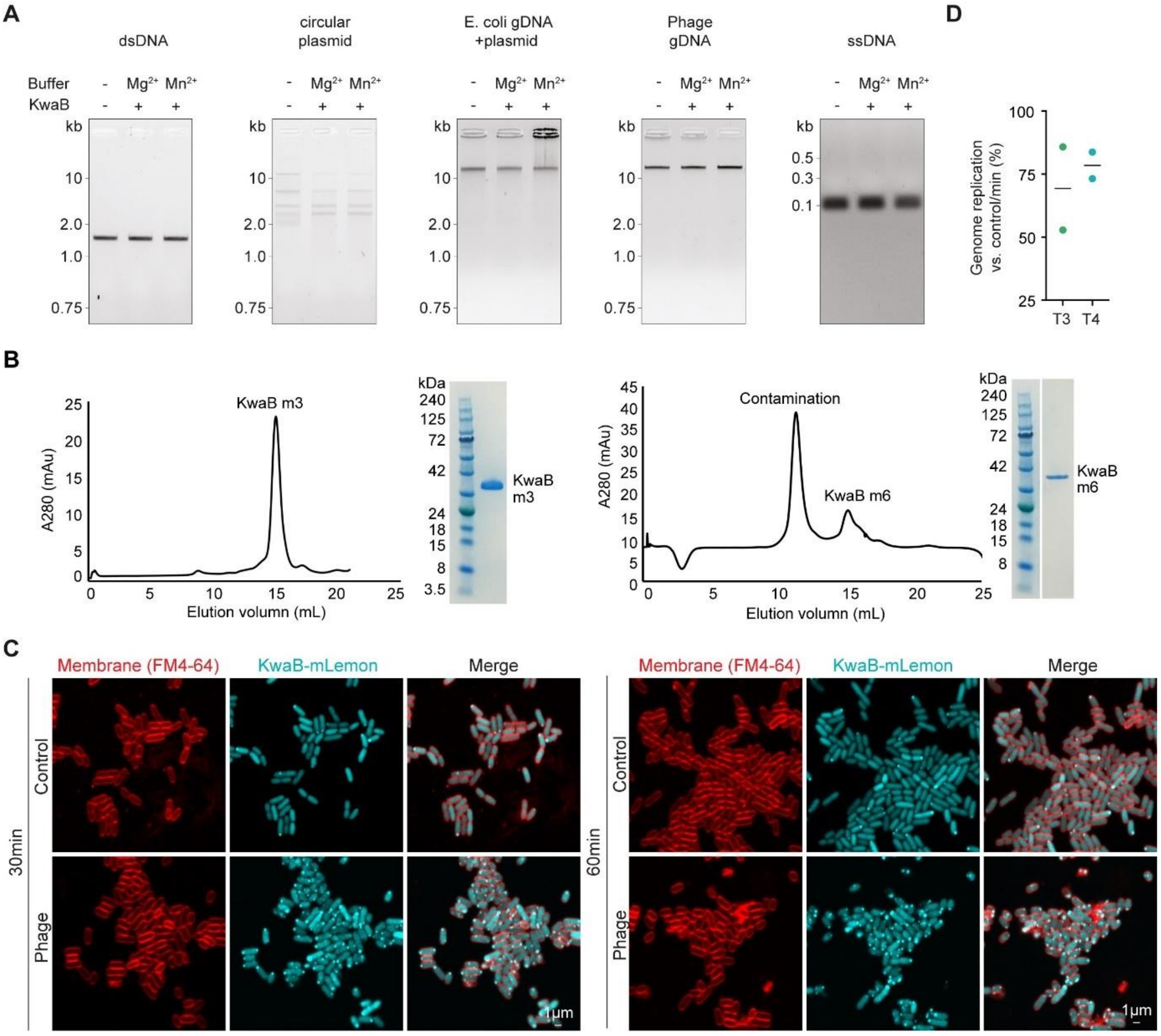


**Figure S6** KwaB does not degrade DNA but binding to phage DNA prevents replication.

**(A)** Analysis of KwaB-mediated nucleic acid cleavage in the presence or absence of $Mg^{2+}$ and $Mn^{2+}$. Reactions contained 500 ng of KwaB and 50 ng of dsDNA or 600 ng ssDNA, and were treated with proteinase K and EDTA before loading on a 1.5% agarose gel.

**(B)** SEC and SDS-PAGE results demonstrate successful expression of KwaB mutants m3 (R142A/R167A/K187A) and m6 (R142A/R167A/K187A/R233A/N134A/D169A).

**(C)** Zoomed-out confocal microscopy images showing KwaB forming membrane-localised foci at 30 and 60 minutes post infection. The cell membrane is stained with FM4-64 (red) and KwaB is tagged with mLemon (depicted in blue).

**(D)** Efficiency of phage genomic DNA (gDNA) replication during infection of control and Kiwa cells. Total DNA was extracted from cells infected with phage at an MOI of 3. The data represents the replication efficiency of phage gDNA per minute over 30 minutes of infection, compared to control cells, and is shown as the mean of two biological replicates with individual data points overlaid.

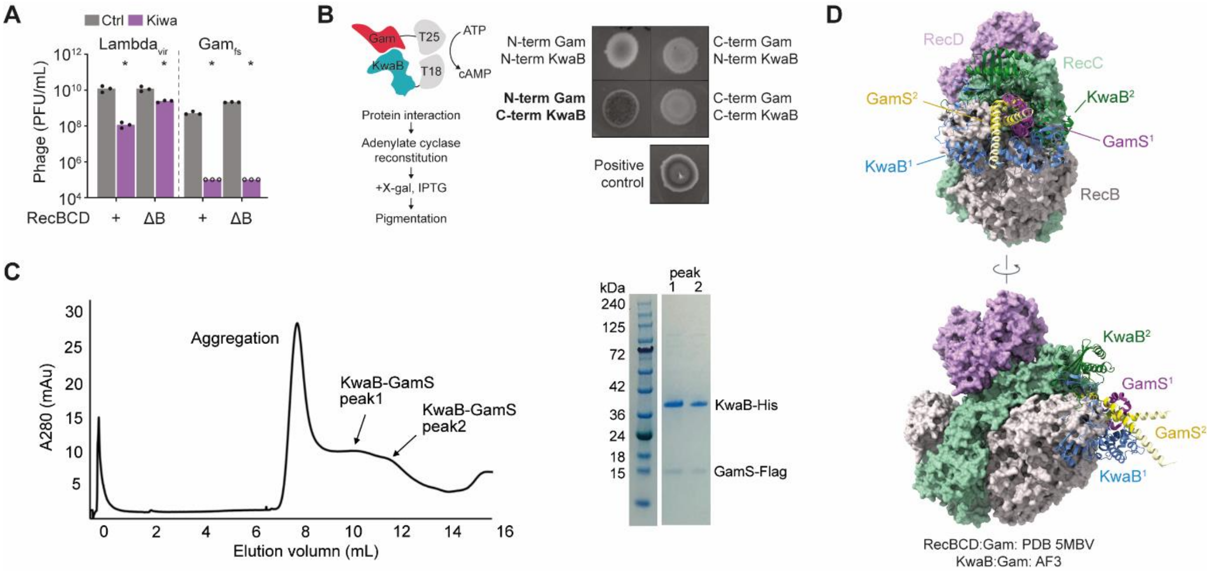


**Figure S7** Inhibition of KwaB by Lambda Gam protein.

**(A)** Kiwa anti-phage activity against phage Lambda$_{vir}$ or a variant with a frameshift mutation in Gam (Gam$_{fr}$). Data represents phage concentration (PFU/ml) on control or Kiwa strains either containing intact RecBCD (+) or a deleted RecB subunit (ΔB). Asterisks indicate a statistically significant decrease of phage concentration relative to control cells on the same host strain (Two-way ANOVA, $p < 0.05$). Bars represent the average of three biological replicates, with individual data points overlaid. Open points indicate instances where it was not possible to determine individual phage plaques, and a value of 1 was assumed at the corresponding dilution.

**(B)** Bacterial two-hybrid analysis of pairwise interactions between constructs of KwaB and Gam fused to T18 and T25 fragments, respectively, and a positive control containing the leucine zipper motif of GCN4.

**(C)** A Flag pull-down assay involving Gam-Flag and KwaB-His proteins, combined with SEC and SDS-PAGE analysis, demonstrated the successful formation of the KwaB-GamS protein complex.

**(D)** Overview of the mutual structural exclusivity of the RecBCD complex inhibited by Lambda Gam (PDB:5MBV) when overlaid with AlphaFold 3 (AF3) predicted KwaB homodimer bound to Gam.
